# Membrane Mimetic-Thermal Proteome Profiling Reveals Broad, Sequence-Independent Membrane Protein Stabilization by Cholesteryl Hemisuccinate

**DOI:** 10.64898/2026.08.17.745344

**Authors:** Ashim Bhattacharya, Shamus Clunie, Frank Antony, Yilun Chen, Hiroyuki Aoki, Mohan Babu, Franck Duong van Hoa

## Abstract

Membrane protein stability is strongly influenced by the surrounding lipid environment, yet how individual lipid species shape membrane proteome stability remains poorly understood. Here, we systematically examined the impact of sphingomyelin, 1,2-dioleoyl-sn-glycero-3-phosphocholine (DOPC), and cholesteryl hemisuccinate (CHS) on membrane proteomes using membrane mimetic platforms combined with membrane mimetic thermal proteome profiling (MM-TPP). CHS shifted the proteome composition away from soluble proteins and toward integral membrane proteins, and induced concentration-dependent thermal stabilization of the mouse liver membrane proteome. Organellar membrane proteins, which displayed greater intrinsic lability than plasma membrane proteins, showed preferential stabilization by CHS. CHS supplementation of *E. coli* membranes similarly produced broad stabilization, indicating that this effect occurs even in cholesterol-naive systems. CHS responses were reproducible across Peptidisc and DDM and independent of CRAC/CARC motif density, supporting a broad, sequence-independent mechanism rather than selective lipid binding, although stabilization was greater among proteins with more transmembrane helices. Accordingly, individual purified proteins reconstituted with CHS exhibited only modest stabilization, consistent with a broad effect that is more apparent at the proteome scale than for any single protein examined in isolation. Together, these findings redefine CHS as a general sterol scaffold that broadly stabilizes membrane proteins and establish MM-TPP as a versatile platform for investigating lipid-dependent effects on membrane proteome stability.

**Subject area:** Integral Membrane Proteins, Thermal Proteome Profiling, Membrane Mimetics, Cholesterol, Lipid-Protein Interactions, Mass Spectrometry

**Highlights:**

- CHS broadly stabilizes membrane proteins across diverse membrane mimetics.
- Organellar membrane proteins exhibit the strongest CHS-mediated stabilization.
- CHS stabilization is conserved in cholesterol-naive *E. coli*.
- MM-TPP enables proteome-wide analysis of lipid-dependent protein stability.

**Graphical Abstract:** 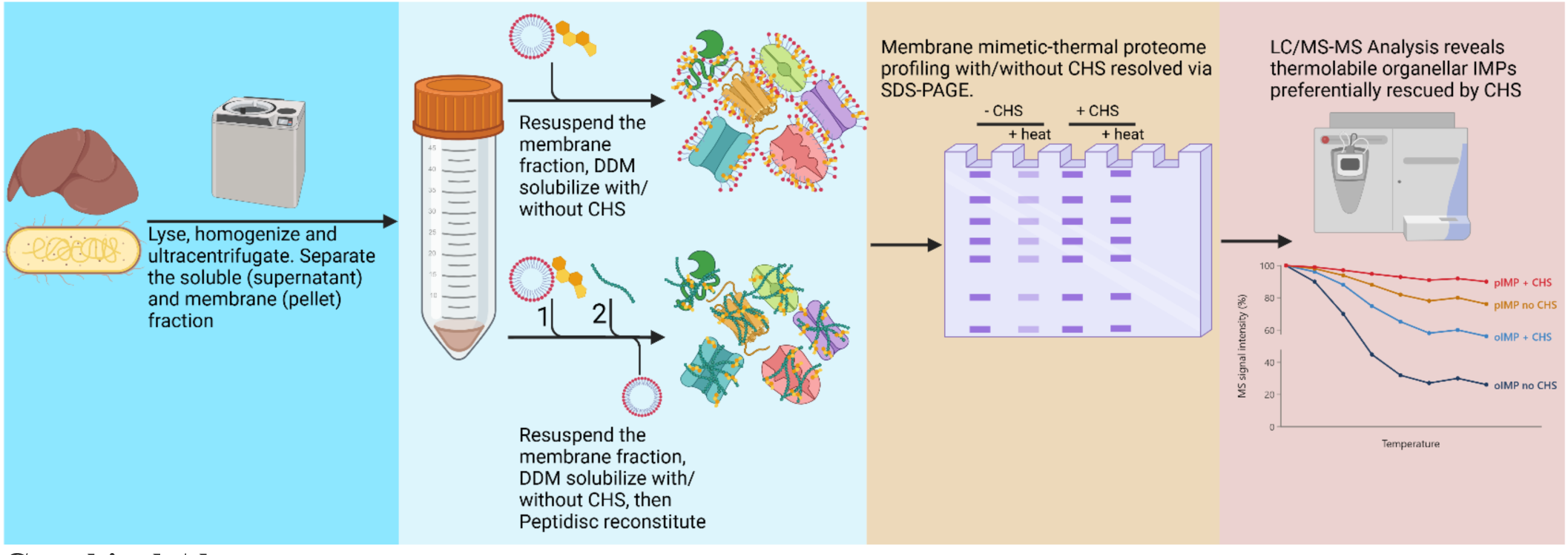

## Introduction

Cholesterol is an essential component of mammalian membranes, where it can comprise up to 50% of total lipid molecules and is differentially distributed across cellular compartments, present at low abundance in organellar membranes and enriched at the plasma membrane, contributing to membrane identity and biophysical diversity^1–3^. In addition to buffering membrane fluidity, cholesterol influences the folding, stability, and function of integral membrane proteins (IMPs) through both indirect effects on membrane physical properties and direct lipid–protein interactions^4^. Structural and biochemical studies have provided evidence of cholesterol-binding sites in receptors, transporters, and ion channels, where cholesterol can influence transmembrane helix packing and conformational transitions^5–8^. Despite its fundamental importance, the extent to which cholesterol contributes to IMP structure, function, and higher-order membrane organization at a systems level remains poorly understood, owing to the pleiotropic nature of its effects and the limited experimental approaches available for disentangling direct cholesterol–protein interactions from broader membrane-mediated effects within complex membrane proteomes^9^.

While cholesterol is the predominant sterol in mammalian membranes, membrane biophysical properties arise from the collective contributions of diverse lipid species that define bilayer order, dynamics, and protein compatibility^10^. Sphingomyelin promotes the formation of ordered lipid raft microdomains through favorable interactions with cholesterol^11^, whereas glycerophospholipids such as 1,2-dioleoyl-sn-glycero-3-phosphocholine (DOPC) favor more fluid, disordered bilayers due to less efficient acyl-chain packing^12^. Cholesteryl hemisuccinate (CHS), a negatively charged cholesterol analog with greater aqueous solubility, serves as a practical cholesterol substitute and is widely used in IMP studies, particularly for GPCRs, because of its reported stabilizing effects on receptor preparations^13,14^.

Together, sphingomyelin, DOPC, and CHS represent lipids with distinct effects on membrane architecture, making them ideal for systematically investigating lipid-dependent effects on the membrane proteome using membrane mimetic platforms. Membrane mimetics are water-soluble systems designed to stabilize IMPs outside the lipid bilayer, including detergents, nanodiscs, and Peptidiscs^15^. While detergents solubilize IMPs through micelle formation, nanodiscs reconstitute IMPs within a defined lipid environment whose composition can be tailored experimentally, enabling the controlled incorporation of specific lipid species^16^. The Peptidisc, by contrast, stabilizes IMPs through a short amphipathic peptide that wraps around transmembrane segments, providing a detergent-free, MS-compatible scaffold that accommodates diverse membrane protein classes^15,17–21^. Furthermore, Peptidisc reconstitution can preserve endogenous annular lipids co-purified with the protein, and exogenous lipids can also be reintroduced to fine-tune the protein microenvironment, with demonstrated effects on protein stability, activity, and protein–protein interactions^22^. This flexibility makes Peptidisc particularly well suited for membrane mimetic-thermal proteome profiling (MM-TPP), a systems-level approach in which IMPs are exposed to a temperature gradient and thermostable fractions are subsequently quantified by liquid chromatography–tandem mass spectrometry (LC-MS/MS)^23^. Differences in thermal dropout rates between conditions provide a proteome-wide measure of changes in protein stability and interaction states^24^.

In this study, we performed lipid-supplemented membrane proteome profiling in Peptidisc- and nanodisc-reconstituted mouse liver membranes to investigate how lipids influence IMP capture and thermal stability. Whereas sphingomyelin and DOPC had minimal effects on membrane proteome recovery, CHS consistently enriched IMPs, prompting further investigation. We therefore applied MM-TPP to mouse liver membrane proteomes in both n-dodecyl-β-D-maltoside (DDM)-solubilized and Peptidisc-reconstituted states. CHS produced a clear dose-dependent thermal stabilization of the membrane proteome, as measured by both SDS-PAGE and LC/MS-MS. Notably, organellar IMPs (oIMPs), which reside in cholesterol-poor membranes and exhibited greater baseline thermal lability than plasma membrane IMPs (pIMPs), showed preferential stabilization in response to CHS. We extended this analysis to *E. coli* membranes as a cholesterol-naive system, where CHS likewise conferred broad stabilization of the membrane proteome. Collectively, this work provides a global view of how lipid supplementation influences membrane proteome thermal stability, while establishing MM-TPP as a platform for the proteome-wide discovery of lipid-dependent effects on IMP composition and stability.

## Results

### CHS Supplementation Enriches IMPs in Peptidisc and spNW30 Libraries

To evaluate the impact of sphingomyelin, DOPC, and CHS supplementation on the capture and stability of the mouse liver membrane proteome, crude membrane fractions from C57BL/6J mice were solubilized with 1% DDM in the presence or absence of 500 µg of each lipid (0.05% w/v), in line with concentrations used for IMP solubilization with CHS in the literature^25,26^. Following solubilization, DDM was diluted to sub-micellar concentrations and exchanged into either His-Peptidisc or His-spNW30 to isolate IMPs into water-soluble assemblies (**Supplemental Figure 1**). These two platforms were previously shown to capture the membrane proteome to comparable depth^15^. Reconstituted assemblies were then affinity purified via their histidine tags to generate an enriched membrane proteome library.

Following reconstitution, the libraries were digested with trypsin and analyzed by bottom-up LC-MS/MS. Identified proteins were categorized by subcellular localization into soluble proteins (Sol), membrane-associated proteins (MAPs), plasma membrane IMPs (pIMPs), and organellar (oIMPs), with pIMPs and oIMPs together defined as total IMPs (tIMPs). Across Peptidisc conditions, lipid supplementation had little effect on overall protein class distribution, with tIMPs consistently comprising ∼40% of total identifications. A modest shift was observed in spNW30 upon CHS supplementation, where tIMP representation increased from 34% to 37% (**Table 1**).

**Table 1.** Number and classification of proteins identified in the mouse liver Peptidisc libraries as a function of the indicated lipid additives. Mouse liver membrane proteins were solubilized in DDM in the presence of 500 µg of the indicated lipid additive (CHS, DOPC, or SM) or in the absence of any lipid supplement, and reconstituted into either Peptidisc or spNW30 nanodiscs prior to MS analysis. For each condition, the total number of detected protein groups (TP) is reported alongside the count and percentage of proteins assigned to each localization category: soluble proteins (SP), membrane-associated proteins (MAP), total integral membrane proteins (tIMP, comprising oIMP and pIMP), organellar integral membrane proteins (oIMP), and plasma membrane integral membrane proteins (pIMP). Percentages are calculated relative to the total detected protein count for that condition. Protein localization assignments and tabulation were performed using dplyr and rendered with knitr::kable. DOPC, dioleoylphosphatidylcholine; CHS, cholesteryl hemisuccinate; SM, sphingomyelin.

|  | TP | TPs |  |  | tIMPs |  |
| --- | --- | --- | --- | --- | --- | --- |
|  |  | SP | MAP | tIMP | oIMP | pIMP |
| Peptidisc | 1033 | 350 (34%) | 257 (25%) | 426 (41%) | 286 (28%) | 140 (13%) |
| + DOPC | 1224 | 437 (36%) | 311 (25%) | 476 (39%) | 316 (26%) | 160 (13%) |
| + CHS | 1324 | 456 (35%) | 336 (25%) | 532 (40%) | 360 (27%) | 172 (13%) |
| + SM | 1225 | 450 (37%) | 312 (25%) | 463 (38%) | 312 (26%) | 151 (12%) |
| spNW30 | 1298 | 517 (40%) | 340 (26%) | 441 (34%) | 307 (24%) | 134 (10%) |
| + DOPC | 1434 | 569 (40%) | 382 (27%) | 483 (33%) | 325 (22%) | 158 (11%) |
| + CHS | 1392 | 530 (38%) | 353 (25%) | 509 (37%) | 337 (25%) | 172 (12%) |
| + SM | 1477 | 593 (40%) | 399 (27%) | 485 (33%) | 321 (22%) | 164 (11%) |

To assess proteome-wide variance across conditions, we performed principal component analysis (PCA) on protein intensities across all samples. The first group (PC1; 35.2%) separated samples by membrane-mimetic platform (Peptidisc versus spNW30), indicating that scaffold identity is the dominant driver of proteome variation. The second group (PC2; 13.5%) separated CHS-supplemented samples from all other conditions, indicating a distinct lipid-dependent shift in the captured proteome across both platforms (**Figure 1A**).

**Figure 1.**
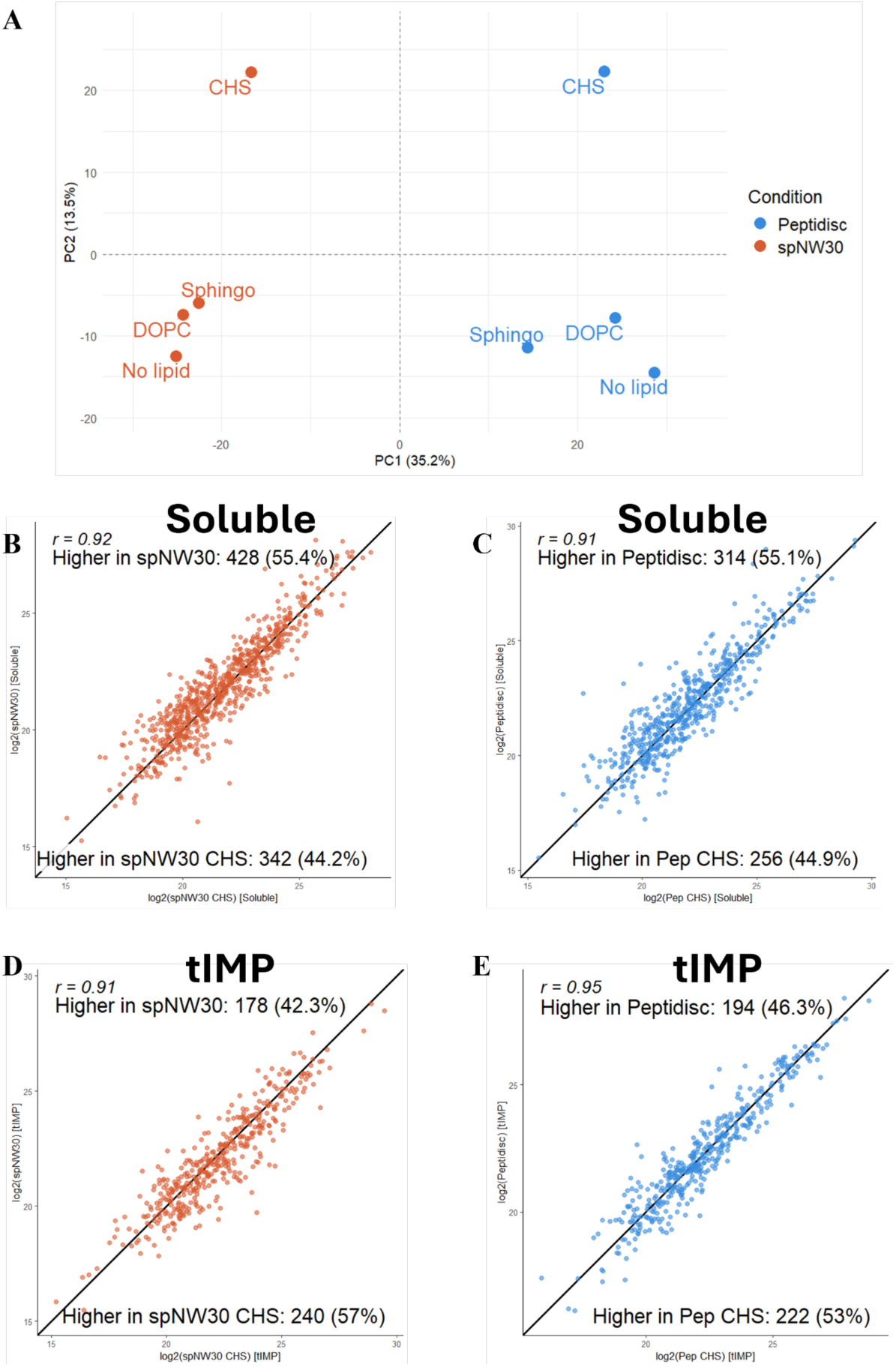
CHS Supplementation Shifts the Mouse Liver Membrane Proteome Toward Integral Membrane Protein Enrichment in Both Peptidisc and spNW30 Nanodiscs. Mouse liver membrane proteome libraries were reconstituted in spNW30 nanodiscs or Peptidisc following DDM solubilization in the presence of 500 µg of the indicated lipid additive (CHS, DOPC, or sphingomyelin). (A) Principal component analysis (PCA) of log2-transformed protein intensities across all conditions, computed using prcomp (R base) and visualized with ggplot2 and ggrepel. (B–E) Pairwise scatter plots comparing log2-transformed protein intensities between the CHS-supplemented and no-additive conditions for soluble proteins (Soluble; B, C) and total integral membrane proteins (tIMP; D, E) identified in each membrane mimetic system. Points above the diagonal indicate proteins more abundant in the no-additive condition; points below indicate proteins enriched in the CHS condition. The proportion of proteins in each direction is indicated as a percentage of all co-detected proteins. Pearson correlation coefficients (r) are shown. Plots were generated using ggplot2.

To determine how CHS altered the capture of protein classes, we compared protein intensity in the CHS-supplemented and unsupplemented conditions across both Peptidisc and nanodisc platforms. Soluble proteins were preferentially enriched in the unsupplemented condition (55% without CHS), whereas tIMPs showed the opposite trend, with 53–57% exhibiting higher intensities under the CHS condition (**Figures 1B–E**). These divergences in the overall protein intensity indicate that CHS preferentially enriches membrane-embedded proteins at the level of protein abundance rather than driving an increase in the number of identified IMPs.

### CHS Preferentially Stabilizes Organellar Integral Membrane Proteins in the Peptidisc Mouse Liver Membrane Proteome

Having established that CHS supplementation alters the captured membrane proteome at the level of increased protein intensity, we next applied MM-TPP to determine whether this enrichment is accompanied by changes in IMP thermal stability and to identify differential dropout rates across the membrane proteome. DDM-solubilized mouse liver membrane fractions were reconstituted into Peptidisc in the presence of 0, 0.5, 1, or 2 mg CHS (0-0.2% w/v) without affinity purification to preserve the native complexity of the membrane proteome. Reconstituted samples were then incubated for 3 min at each of five discrete temperatures (50, 60, 70, 80, and 100°C) to generate a thermal denaturation series. Thermostable fractions were recovered by ultracentrifugation and analyzed by SDS-PAGE (**Figure 2A**). Inspection and densitometric analysis of the gels revealed a dose-dependent stabilization effect, with samples supplemented with 2 mg CHS retaining 67% of the room-temperature signal at 80°C compared to 49% in the unsupplemented condition (**Figure 2B**).

**Figure 2.**
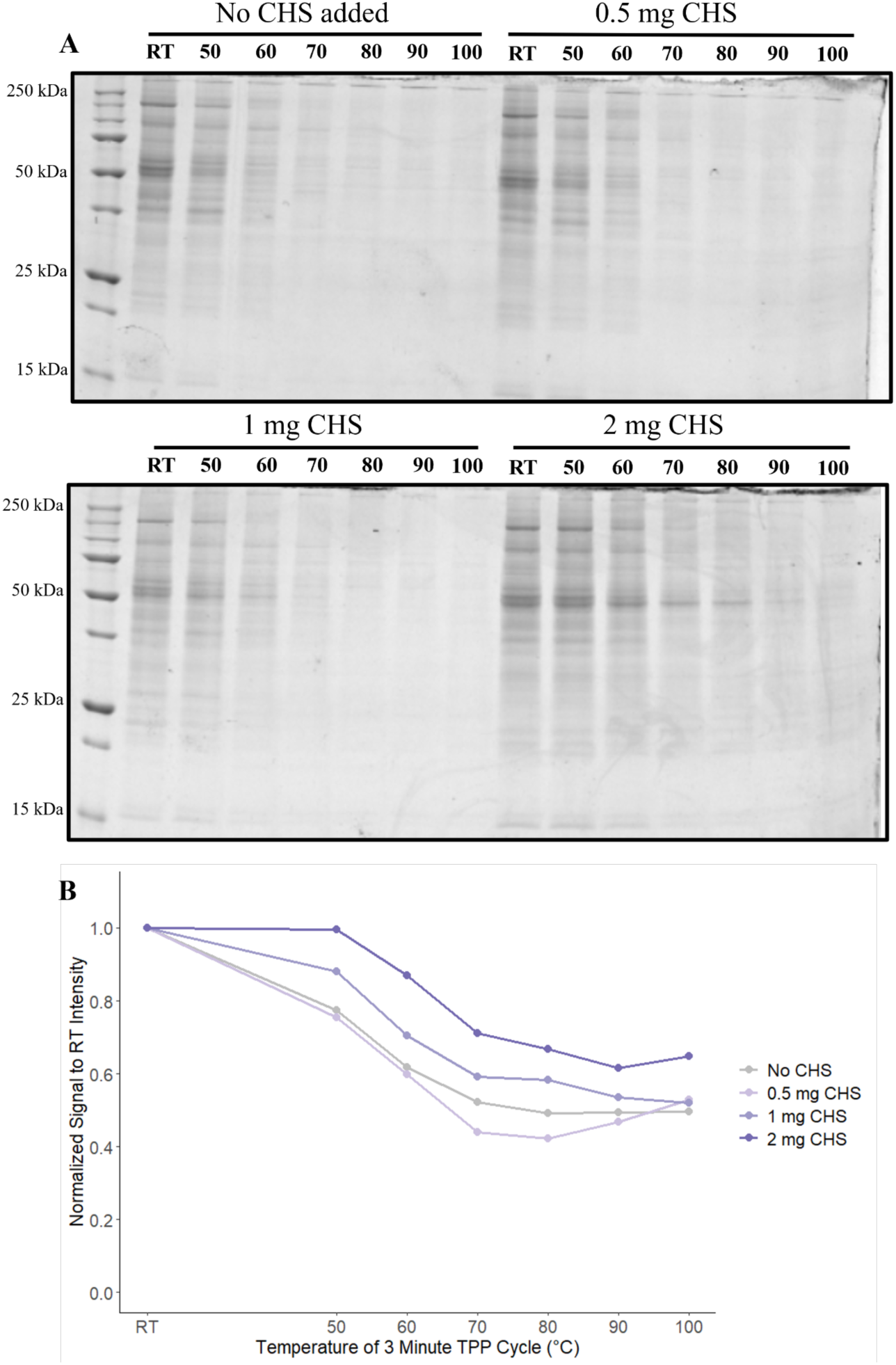
CHS Supplementation Confers Dose-Dependent Thermal Stabilization of the Mouse Liver Peptidisc Membrane Proteome. Mouse liver membrane proteomes were reconstituted in Peptidisc following DDM solubilization in the presence of 0, 0.5, 1, or 2 mg CHS. (**A**) SDS-PAGE analysis of the soluble fraction after 3-minute thermal treatment at the indicated temperatures (RT–100°C), with molecular weight markers indicated on the left (kDa). **(B)** Thermal retention curves derived from densitometric quantification of Peptidisc fractions shown in (A). Total lane intensity between 250 and 15 kDa was normalized to the RT value for each condition to generate a retention curve. Densitometry was performed using ImageJ, and retention curves were plotted using ggplot2.

To resolve changes associated with this response at the protein level, the CHS supplemented (2mg) and unsupplemented fractions were analyzed by LC-MS/MS, with protein ID counts summarized in **Supplemental Table 1**. Across all temperatures, the CHS-supplemented condition identified more total proteins than the unsupplemented condition (e.g., 1775 vs. 1613 at RT; 1140 vs. 668 at 100°C). The CHS-supplemented condition also retained more proteins overall at the highest temperature, with 64.2% of the starting proteome still detected at 100°C, compared to only 41.4% in the unsupplemented condition. Additionally, the CHS-supplemented condition showed a higher proportion of oIMPs across all temperatures (15– 17%) relative to the unsupplemented condition (10–15%).

For each protein, retention was calculated as the ratio of MaxLFQ intensity at each temperature point relative to the room temperature sample, generating a thermal retention curve across the denaturation series. Mean retention curves were then computed across all oIMPs and pIMPs in the 0 and 2 mg CHS conditions to compare class-level thermal behavior. In the absence of CHS, oIMPs showed greater thermal signal loss than pIMPs, with a mean retention difference of 31% when averaging normalized intensities across all temperature points. With CHS, a preferential stabilization of oIMPs relative to pIMPs was obtained across the temperature series, with mean increases in abundance of 28% and 17%, respectively (**Supplemental Figure 2A**). To further evaluate the distribution of CHS-induced stabilization, protein intensities at 100°C were directly compared between the 0 and 2 mg CHS conditions for each oIMPs and pIMPs using scatter plots. This analysis further supported the retention trend observed, with 89% of oIMPs and 66% of pIMPs showing higher abundance in the 2 mg CHS condition at 100°C relative to the unsupplemented control (**Supplemental Figure 2B–C**). Together, these data indicate a preferential stabilizing effect of CHS on oIMPs within the Peptidisc-reconstituted membrane proteome under MM-TPP conditions.

### CHS Confers Concentration-Dependent and Preferential Thermal Stabilization of oIMPs in DDM-Solubilized Mouse Liver Membranes

The initial characterization of CHS-induced IMP stabilization was performed in the Peptidisc and nanodisc system; however, the majority of CHS-supplemented IMP solubilization studies in the literature employ DDM detergent directly. To determine whether the stabilizing effect of CHS extends to this more widely used context and to maximize the translational relevance of our findings, we assessed whether CHS-induced stabilization was retained in DDM-solubilized membrane preparations. Mouse liver crude membranes were solubilized with 1% DDM in the presence of 0, 1, 2, or 4 mg CHS (0-0.4% w/v) and subjected to thermal denaturation at 51, 60, 70, 80, and 90°C for 3 minutes. Thermostable fractions were recovered by ultracentrifugation and analyzed by SDS-PAGE (**Figure 3A**). At 80°C, total protein retention increased in a CHS-dependent manner (25%, 39%, 48%, and 59% for 0–4 mg CHS, respectively), as determined by whole-lane densitometric analysis normalized to the room temperature control, demonstrating concentration-dependent stabilization of the DDM-solubilized proteome **(Figure 3B**).

**Figure 3.**
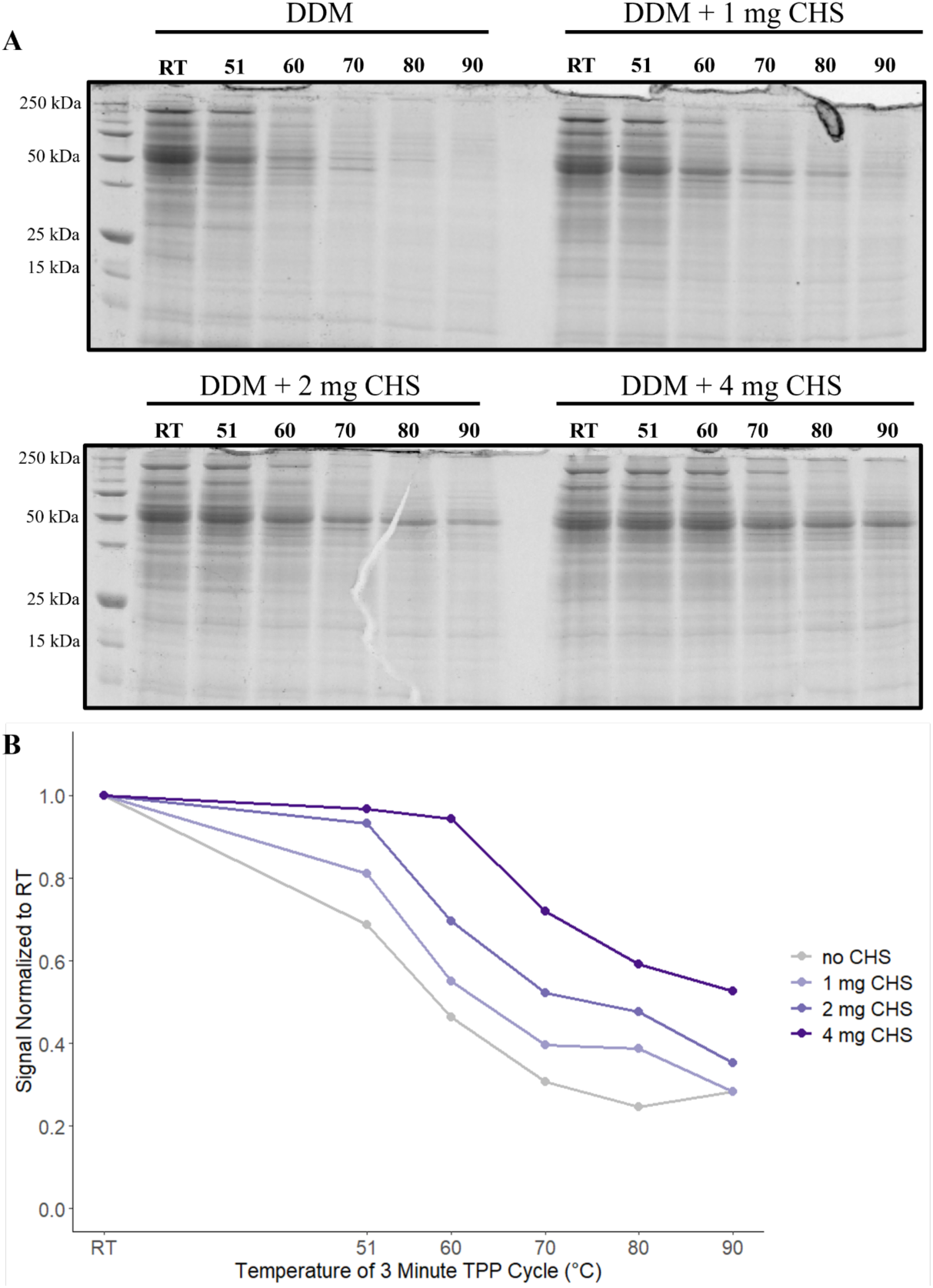
CHS Supplementation Confers Dose-Dependent Thermal Stabilization of the Mouse Liver DDM-Solubilized Membrane Proteome. The mouse liver crude membrane was solubilized in DDM supplemented with 0, 1, 2, or 4 mg CHS and subjected to thermal incubation for 3-minutes at the indicated temperatures (RT–90°C), followed by ultracentrifugation to pellet aggregated proteins. (**A**) SDS-PAGE analysis of the supernatant fractions Molecular weight markers are indicated on the left (kDa). (**B**) Thermal retention curves derived from densitometric quantification of the DDM fractions shown in (A). Total lane intensity was normalized to the RT value for each condition to generate a retention curve. Densitometry was performed using ImageJ, and retention curves were plotted using ggplot2.

These samples were processed by SP4 cleanup^15^ and analyzed by LC-MS/MS with protein ID counts found in **Supplemental table 2**. The 4 mg CHS-supplemented condition retained more proteins overall at the highest temperature, with 91.8% of the starting proteome still detected at 90°C, compared to only 64.2% in the unsupplemented DDM condition. Additionally, the CHS-supplemented condition maintained a more stable proportion of oIMPs across the thermal range (13–15%), whereas the DDM condition showed a steady decline (15% down to 11%).

The MaxLFQ intensities were normalized for each protein at each temperature to the room temperature control to generate per-protein retention profiles across the denaturation series. Mean retention was then computed across all quantified oIMPs and pIMPs separately for the 0 and 4 mg CHS conditions. In the absence of CHS, oIMPs showed greater thermal signal loss than pIMPs, with a mean difference of 33% in normalized retention values averaged across all temperature points of the gradient. The CHS supplementation reversed this trend, with oIMPs displaying a mean increase of +28% in normalized retention relative to the unsupplemented condition, compared to +2% for pIMPs (**Supplemental Figure 3A**). This pattern was corroborated by scatter plot comparison of MaxLFQ intensities at 90°C between the 0 and 4 mg CHS conditions, where 70.9% of oIMPs were enriched in the CHS condition compared to only 50.4% of pIMPs (**Supplemental Figure 3B-C**), reinforcing that the stabilizing effect of CHS is preferential to oIMPs rather than a uniform increase across all IMP classes. Together, these results confirm that CHS preferentially stabilizes oIMPs in DDM-solubilized membrane preparations, consistent with the Peptidisc findings and supporting a broadly conserved stabilizing effect that is independent of the membrane mimetic or reconstitution platform.

### CHS-Induced Stabilization Is Reproducible Across Membrane Mimetic Systems and Reflects a Broad, Motif-Independent Mechanism Modulated by Localization-Specific Structural Features

Having established that CHS enriches tIMPs in both DDM and Peptidisc preparations, we next asked whether CHS-induced effects are reproducible across membrane mimetic systems and whether the magnitude of stabilization is influenced by protein localization or structural features. To address these questions, we compared CHS-induced abundance changes between Peptidisc and DDM datasets and examined potential relationships between thermal stabilization and protein localization, cholesterol-recognition motifs, transmembrane topology, and TM-segment abundance.

PCA of all samples showed that PC1 (39.4%) separated the two membrane mimetic systems, while PC2 (12.9%) captured temperature-dependent variation, with heated samples converging across conditions (**Figure 4A**). Comparison of per-protein CHS-induced log2 fold changes correlated more strongly between Peptdisic and DDM for oIMPs (r = 0.55) and pIMPs (r = 0.54) than for soluble proteins (r = 0.39) or MAPs (r = 0.31) (**Figure 4B–E**), indicating that CHS responses are most reproducible among IMPs.

**Figure 4.**
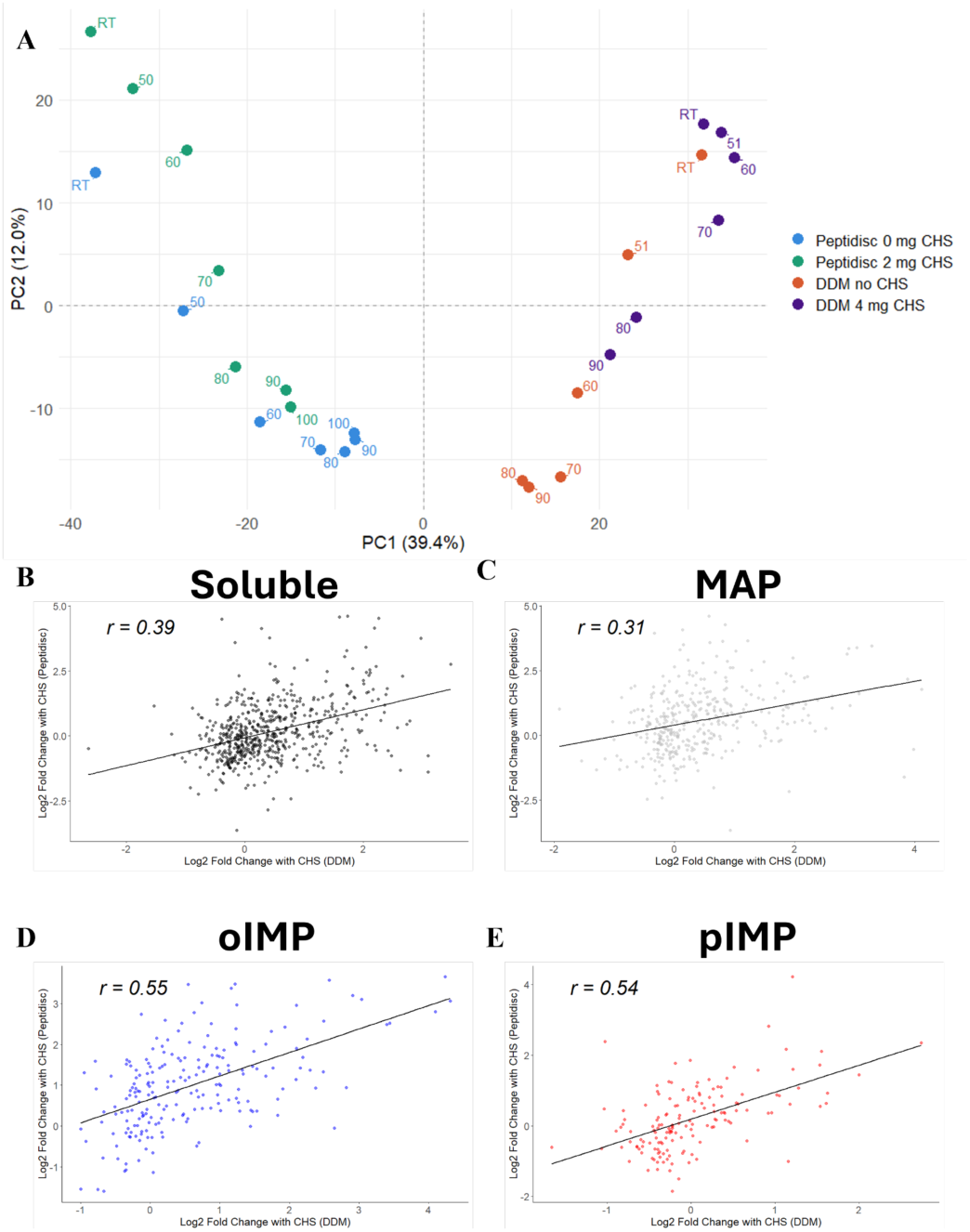
Comparison of CHS-Induced Protein Abundance Changes in DDM and Peptidisc Membrane Proteomes by Membrane Protein Class. Mouse liver membrane proteins were solubilized in DDM with or without 4 mg CHS and analyzed by DDA-MS, or independently solubilized and reconstituted into Peptidisc with 0 or 2 mg CHS and analyzed by DIA-MS. **(A)** Principal component analysis of log2-transformed protein intensities across all DDA and DIA replicate samples, restricted to proteins detected in both datasets. Missing values were imputed by per-protein column median, and genes with zero variance across samples were excluded prior to PCA. PCA was computed using prcomp (centered and scaled) and visualized with ggplot2 and ggrepel. Each point represents one replicate injection, colored by dataset and CHS condition (DDM no CHS, DDM 4 mg CHS, Peptidisc 0 mg CHS, Peptidisc 2 mg CHS) and labeled by temperature to illustrate the TPP trajectory of each condition through PCA space. **(B–E)** Pairwise scatter plots comparing the log2 fold change conferred by CHS in DDM (x-axis) versus Peptidisc (y-axis), stratified by protein localization category: soluble (B, black), membrane-associated proteins (C, grey), organellar integral membrane proteins (D, blue), and plasma integral membrane proteins (E, red). The x-axis represents the mean log2 intensity difference between the 4 mg CHS and no CHS DDM conditions; the y-axis represents the mean log2 intensity difference between the 2 mg CHS and 0 mg CHS Peptidisc conditions. Only proteins co-detected in both datasets were included, joined by gene name using inner_join from dplyr. A linear regression line is overlaid and Pearson correlation coefficients (r) are shown. Plots were generated using ggplot2.

We next examined whether CHS-induced thermal stabilization could be explained by specific cholesterol-recognition motifs or IMP architecture, using a retention-based stabilization score (**Table 2**). CRAC/CARC motif density^27,28^ did not significantly predict stabilization in either dataset, and stabilization did not differ between α-helical and β-barrel IMPs in DDM nor Peptidisc. In contrast, TM helix abundance correlated significantly with stabilization in both datasets, with proteins containing more transmembrane helices showing greater CHS-induced stabilization. These results suggest that CHS-mediated stabilization does not depend on canonical cholesterol-recognition motifs or on α-helical versus β-barrel architecture, though its magnitude is modulated by transmembrane helix content.

**Table 2:** Association of Protein Localization and Structural Features with CHS-Induced Thermal Responses. Linear models, Spearman correlations, and Wilcoxon rank-sum tests were performed across Peptidisc and DDM datasets to assess whether CRAC/CARC motif density within transmembrane segments, transmembrane helix count, or helical versus beta-barrel topology predict CHS-induced stabilization. Stabilization was quantified as a retention-based Stabilization Score (mean Δretention across all non-RT temperatures between CHS and no-CHS conditions). CRAC (cholesterol recognition amino acid consensus) and CARC motifs are cholesterol-binding sequence motifs found within transmembrane domains of integral membrane proteins. CRAC/CARC motif density and transmembrane helix counts were derived from Phobius-predicted transmembrane topology annotations and UniProt protein sequences retrieved via the UniProt REST API. Helical versus beta-barrel topology classification was obtained from UniProt feature annotations queried using the jsonlite and httr packages. All statistical analyses were performed in R using the tidyverse, rstatix, and base stats packages. N refers to the number of proteins detected in both CHS and no-CHS conditions with a calculable Stabilization Score. The helical versus beta-barrel comparison was performed for the DDM dataset only, as the Peptidisc dataset contained insufficient beta-barrel representation for statistical testing.

|  | Analysis | Test | Statistic | P-value | N |
| --- | --- | --- | --- | --- | --- |
| Peptidisc | CRAC/CARC motif abundance | Linear model | $R^2 = 0.00115$ | 0.794 | 402 |
| | TM helix abundance vs. Stabilization Score | Spearman correlation | $\rho = 0.175$ | 0.000433 | 402 |
| | Helical vs. beta-barrel TM topology | Wilcoxon rank-sum | $W = 1282$ | 0.767 | 399<br>(Helical n=392, Beta-barrel n=7) |
| DDM | CRAC/CARC motif abundance | Linear model | $R^2 = 0.00807$ | 0.0766 | 637 |
| | TM helix abundance vs. Stabilization Score | Spearman correlation | $\rho = 0.135$ | 0.000666 | 637 |
| | Helical vs. beta-barrel TM topology | Wilcoxon rank-sum | $W = 3116$ | 0.231 | 633<br>(Helical n=625, Beta-barrel n=8) |

To assess whether the magnitude of CHS-induced rescue varies by localization class, we compared a per-protein CHS-induced retention rescue score (mean retention normalized to RT intensity, with CHS minus without CHS, averaged across the TPP series) across Soluble, MAP, oIMP, and pIMP proteins. This revealed a significant overall difference in DDM but not in Peptidisc (**Supplemental Figure 4**). Pairwise comparisons within DDM showed that pIMP proteins had significantly less rescue than oIMP, MAP, and Soluble proteins, while oIMP showed greater rescue than Soluble; oIMP/MAP and Soluble/MAP did not differ. No corresponding contrasts were significant in Peptidisc (**Supplemental Figure 4**).

In DDM, both baseline thermal lability and CHS-induced rescue varied significantly among organellar subclasses, with mitochondrial oIMPs representing the most thermolabile population and receiving the greatest rescue (**Supplemental Figure 5**). We next examined TM helix count within pIMPs and oIMPs separately. TM segment count was significantly higher in pIMPs than oIMPs in Peptidisc but not DDM (**Supplemental Table 3**). Independently of this difference in TM abundance between classes, CHS-induced rescue itself grew with increasing TM-segment count among pIMPs in both Peptidisc and DDM, whereas no corresponding relationship was observed for oIMPs in either dataset (**Figure 5**). Together, these findings show that CHS-induced rescue, while broad and motif-independent, is shaped by organelle identity in oIMPs and TM helix count in pIMPs.

**Figure 5.**
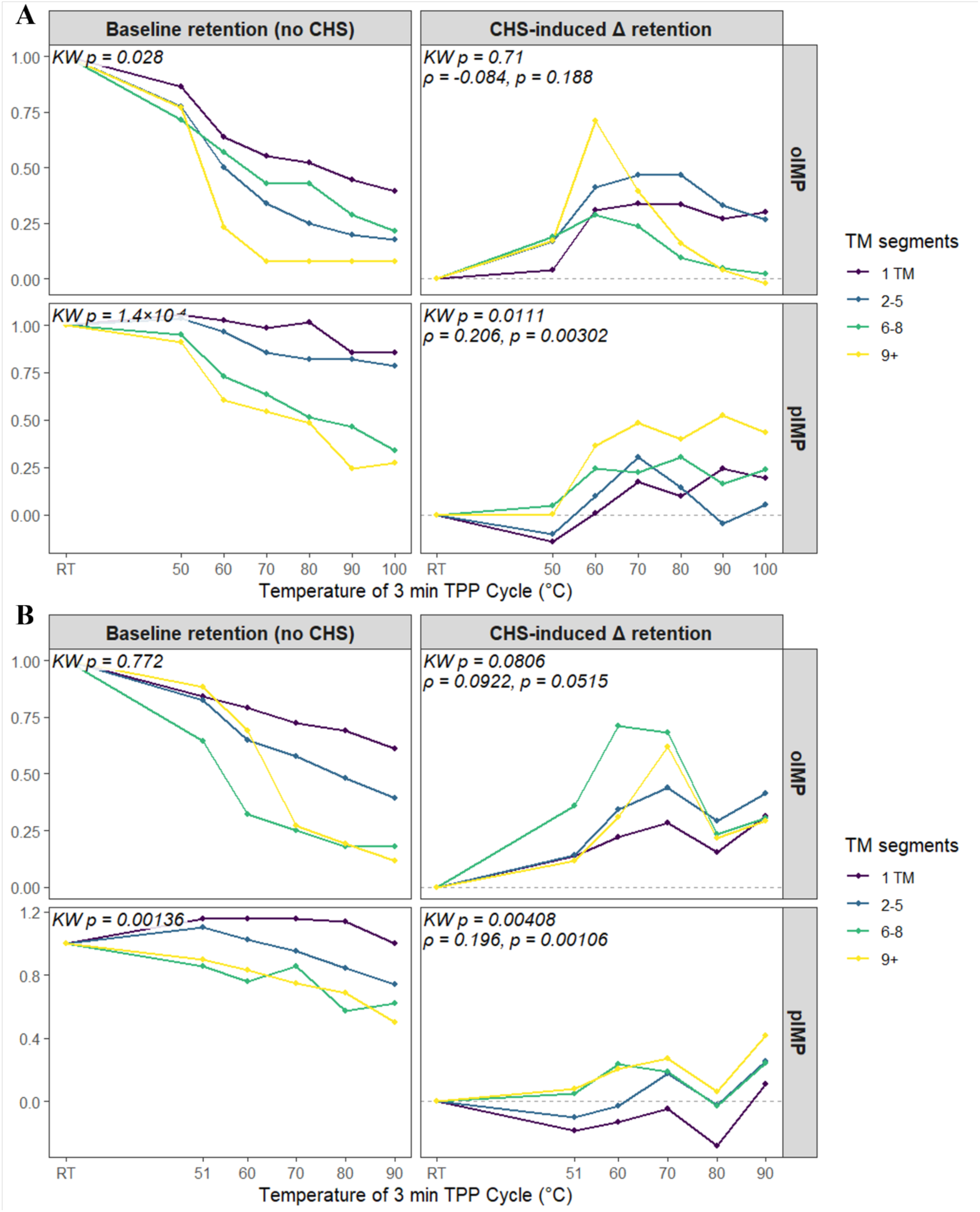
Baseline thermal lability scales with transmembrane segment count, and CHS-induced rescue correlates with TM abundance in pIMPs but not oIMPs, across Peptidisc and DDM datasets. oIMP and pIMP proteins were binned by transmembrane (TM) segment count (1 TM, 2–5, 6–8, 9+) using phobius and retention curves were generated across the TPP temperature series for Peptidisc and DDM, separately for baseline retention (no CHS, left) and CHS-induced Δ retention (right). **(A)** In Peptidisc, baseline retention differed significantly by TM bin in both oIMP (Kruskal-Wallis p = 0.028) and pIMP (p = 1.4×10⁻⁴). CHS-induced Δ retention did not correlate with TM bin in oIMP (KW p = 0.71; Spearman ρ = −0.084, p = 0.188), but showed a significant positive correlation in pIMP (KW p = 0.0111; ρ = 0.206, p = 0.00302). **(B)** In DDM, baseline retention did not differ significantly by TM bin in oIMP (KW p = 0.772) but did in pIMP (KW p = 0.00136). CHS-induced Δ retention showed a trend toward significance in oIMP (KW p = 0.0806; ρ = 0.0922, p = 0.0515) and a significant positive correlation in pIMP (KW p = 0.00408; ρ = 0.196, p = 0.00106). Dashed lines mark zero Δ retention. Statistical analysis was performed in R using kruskal.test and Spearman cor.test, with data processing in dplyr/tidyr and figures generated using ggplot2.

Finally, to exclude an artifact arising from detergent–sterol micelle formation, we compared CHS addition before versus during DDM solubilization. Protein retention at 90°C was similar between conditions (54% vs. 58% of RT signal), arguing against a major contribution from detergent–sterol interactions and supporting direct CHS-dependent stabilization of IMP (**Supplemental Figure 6**).

### CHS-Induced Thermal Stabilization Extends to the Cholesterol-Naive *E. coli* Membrane Proteome

Having characterized CHS-induced thermal stabilization in mouse liver membranes, we next asked whether this effect requires a native cholesterol-containing membrane background, or whether it can occur independently of endogenous cholesterol. To address this, we performed MM-TPP on *E. coli* crude membrane fractions solubilized with 1% DDM, which were either retained in detergent or further reconstituted into Peptidisc, each supplemented with 4 mg CHS (0.4% w/v). Consistent with mouse liver results, SDS-PAGE analysis showed increased protein retention across the full temperature gradient in both preparations (**Figure 6A**), with mean signal intensities increasing by 16% and 24% relative to unsupplemented controls, respectively (**Figure 6B**). Because *E. coli* membranes lack cholesterol, these findings support the notion that CHS-mediated stabilization can occur independently of endogenous cholesterol-dependent interactions and may represent a broadly acting effect.

**Figure 6.**
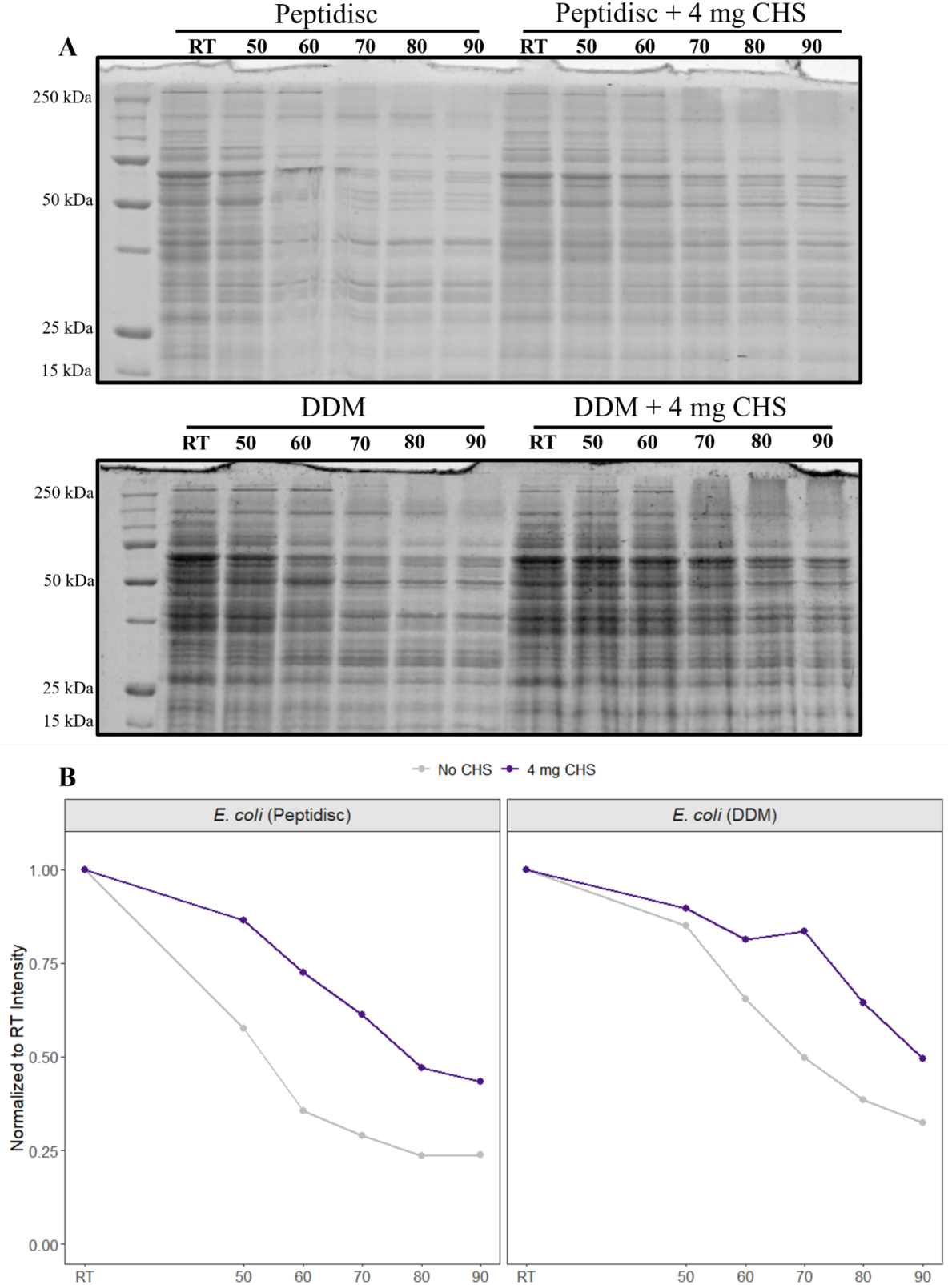
CHS-Mediated Thermal Stabilization Is Conserved in the Cholesterol-Naive *E. coli* Membrane Proteome Across DDM and Peptidisc Platforms. (**A**) SDS-PAGE analysis of Peptidisc reconstituted and DDM-solubilized *E. coli* membrane proteome across the temperature gradient. (**B**) Thermal retention curves derived from densitometric quantification of DDM *E. coli* fractions shown in (A). Total lane intensity between 250 and 15 kDa was normalized to the RT value for each condition to generate a retention curve. Densitometry was performed using ImageJ, and retention curves were plotted using ggplot2.

### CHS-Induced Stabilization Is Detectable but Modest at the Level of Individual Purified E. coli IMPs

Having characterized broad, proteome-wide CHS-induced stabilization, we next asked whether this aggregate effect is reflected in thermal stabilization at the level of individual purified IMPs, which would be of greater practical interest to researchers working with single IMP targets. Three *E. coli* IMPs were selected for recombinant expression and Ni-NTA purification based on expression yield and stability: YidC, MsbA, and a YibN-TMS-APEX2 fusion protein, chosen for their high expression levels suitable for robust TPP densitometry.

Prior to thermal profiling, we first assessed whether CHS supplementation (0.4% w/v) itself altered protein recovery or purity across the membrane mimetic systems used in this study. In Peptidisc-reconstituted preparations, a reduction in recovery was observed (**Supplemental Figure 7**); however, CHS showed no detectable effect on solubilization yield or sample purity for any of the three proteins (**Supplemental Figure 8**). No changes in oligomerization state were observed in Peptidisc or DDM, as assessed by native PAGE under clear and blue-native conditions (**Supplemental Figures 9–11**). Similarly, for MsbA solubilized using the PDET-1 Peptergent system, CHS did not alter extraction efficiency or purity (**Supplemental Figure 12**)^29^. Together, these results indicate that CHS does not broadly perturb protein recovery across membrane mimetics.

We then performed MM-TPP on the purified proteins, using SDS-PAGE densitometry to quantify thermal retention, to assess whether proteome-level stabilization trends are reflected at single-protein resolution. The YibN-TMS-APEX2 fusion exhibited the most pronounced response, with mean increases in thermal retention of 15.6% and 7.6% in DDM and Peptidisc, respectively, relative to no CHS controls (**Figure 7**). YidC showed a smaller but consistent positive effect, with retention increases of 8.9% (DDM) and 6.6% (Peptidisc) **(Supplemental Figure 13**). MsbA showed a modest increase in Peptidisc (3.7%) but no change in DDM (– 0.2%), indicating that CHS did not stabilize MsbA in this condition (**Supplemental Figure 14**). Collectively, these results indicate that CHS-induced stabilization is detectable but modest and protein-dependent at the individual protein level, with some proteins (e.g., MsbA in DDM) showing negligible response, consistent with the proteome-wide effect reflecting the cumulative contribution of many such variable, protein-dependent responses.

**Figure 7.**
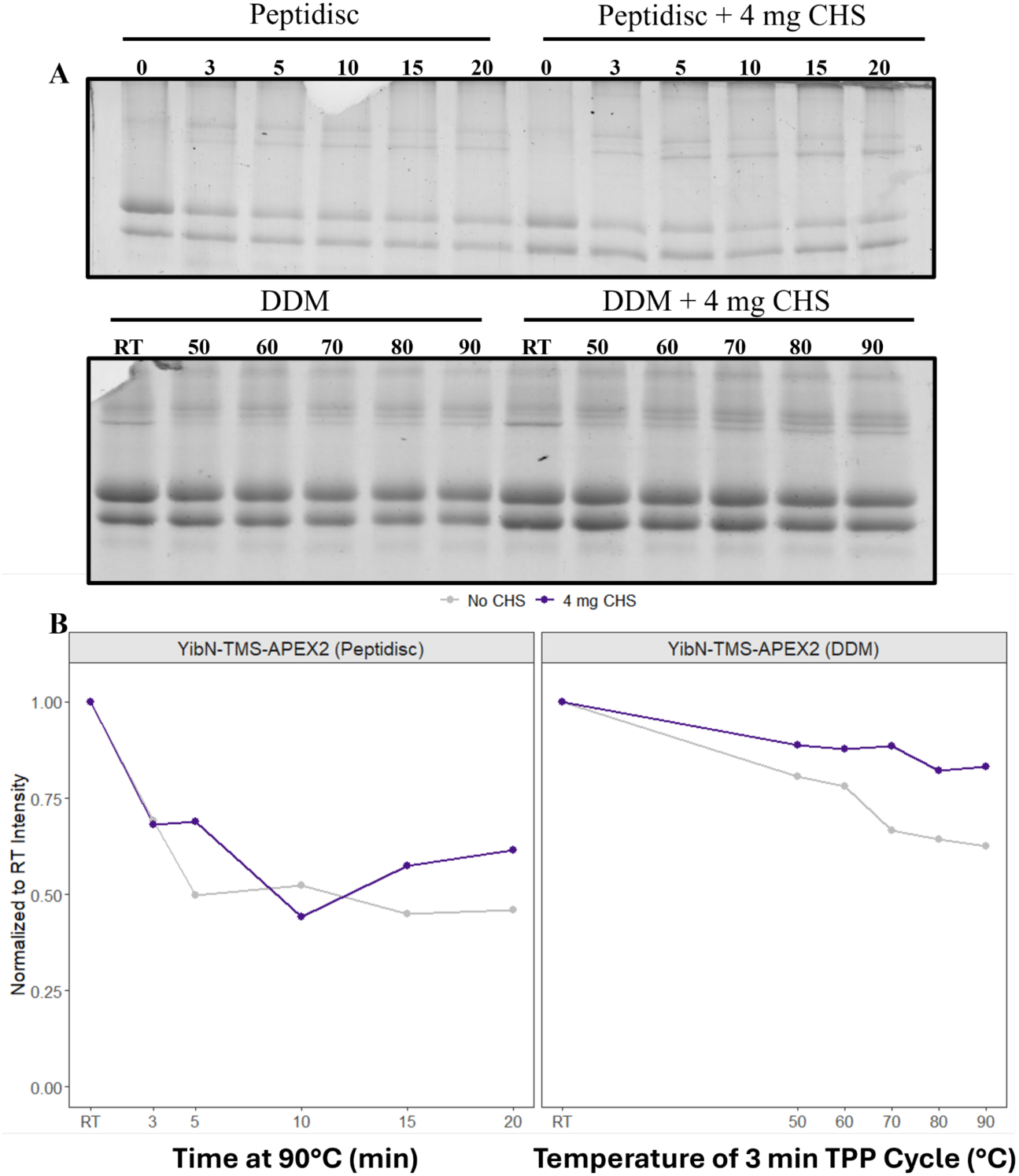
Effect of CHS Supplementation on Thermal Profiles of Purified YibN-TMS-APEX2 in DDM and Peptidisc. **(A)** Peptidisc Reconstituted and DDM-solubilized YibN-TMS-APEX2 supplemented with or without 4 mg CHS were subjected to 3-minute thermal treatment at the indicated temperatures, followed by ultracentrifugation to pellet aggregated protein, and the soluble supernatant was analyzed by SDS-PAGE. Molecular weight markers are indicated in kDa. **(B)** Densitometric quantification of whole-lane signal from the SDS-PAGE gels shown in (A) performed with ImageJ, normalized to the room temperature (RT) or time zero (0 min) lane value for each condition. The two panels retain independent x-axes reflecting their different temperature ranges, rendered using facetted_pos_scales from ggh4x and plotted with ggplot2.

## Discussion

CHS is frequently used as a soluble cholesterol substitute and IMP stabilizer, with concentrations typically ranging from 0.03–0.2% (w/v) in IMP solubilization workflows^25,26,30–32^ and thermal shift assays^33–38^. Accordingly, several studies have reported particular benefits for specific IMP classes, most notably GPCRs^30,39–41^, implicitly attributing stabilization to discrete cholesterol–protein interactions; cholesterol itself has further been described as an allosteric modulator of GPCR activity, enhancing agonist binding and G-protein coupling, with CHS used as its detergent-compatible analog in such assays^42,43^. Whether this benefit holds at the membrane proteome level, however, has not been systematically evaluated. Here, using membrane proteome profiling and MM-TPP across multiple membrane mimetic systems and a CHS concentration range (0.05–0.4% w/v) overlapping with those used in prior workflows, we show that CHS produces broad, concentration-dependent stabilization of IMPs in both mouse liver and cholesterol-naive *E. coli* membranes. The reproducibility of this response across detergent micelles, Peptidisc, and nanodisc assemblies indicates that CHS-mediated stabilization reflects a general effect on IMPs rather than a phenomenon restricted to a particular membrane mimetic or endogenous cholesterol background. Moreover, the independence of stabilization from transmembrane topology, protein architecture and predicted CRAC/CARC motif density argues against a mechanism dominated by specific cholesterol-binding sites.

A notable observation was that organellar IMPs (oIMPs) displayed greater baseline thermal lability than plasma membrane IMPs (pIMPs) across both DDM and Peptidisc datasets, yet also experienced the greatest rescue upon CHS supplementation. To our knowledge, this proteome-wide difference in thermal behavior has not previously been reported. Given that organellar membranes are relatively poor in cholesterol compared with the plasma membrane, we speculate that oIMPs may have subtly different lipid requirements or form less intimate associations with their native lipid environment, potentially resulting in greater lipid loss during extraction and purification. This could contribute to their reduced intrinsic stability and increased sensitivity to sterol supplementation, though further work would be needed to directly test this hypothesis. Consistent with a structural contribution to this rescue, CHS-induced stabilization correlated positively with transmembrane helix abundance across the membrane proteome in both DDM and Peptidisc datasets, an effect that was most pronounced among pIMPs specifically, suggesting that proteins with more extensive TM architecture are, on average, more responsive to CHS supplementation. Thus, while CHS-mediated stabilization is broad and sequence-independent, its magnitude appears to be modulated by localization-specific membrane environments and intrinsic structural features.

The minimal effects of DOPC and sphingomyelin indicate that stabilization is not a general consequence of lipid supplementation, pointing instead to the rigid sterol scaffold of CHS as the basis for this broad stabilization mechanism. Previous studies likewise found that only sterols retaining a cholesterol-like rigid core stabilized GPCRs and membrane transporters despite differences in headgroup chemistry and micelle properties^38,44^. Consistent with these findings, CHS-mediated enrichment persisted across detergent micelles, Peptidisc, and nanodiscs regardless of whether CHS was added before or after detergent solubilization, echoing a prior report of CHS stabilizing a purified IMP added post-purification^45^, thereby arguing against detergent–sterol micelle formation as the primary mechanism and further supporting a direct sterol effect. Specific and nonspecific sterol interactions can coexist, and the broad stabilization observed here is consistent with previous studies reporting nonspecific CHS effects alongside bona fide cholesterol-binding sites within the same protein^46^. Consequently, stabilization within commonly used CHS concentration ranges should not, on its own, be interpreted as evidence of specific cholesterol binding.

These findings have practical implications for experimental workflows that routinely incorporate CHS, including detergent solubilization, cryo-EM sample preparation, crystallography, and thermal stability assays. Because GPCRs are the IMP class most commonly supplemented with CHS, applying MM-TPP to GPCR-rich proteomes such as brain tissue^47^ or recombinant systems, would directly test whether the broad stabilization effect observed here extends to this receptor family. Future work could further define the mechanism of CHS-mediated stabilization through systematic variation of sterol scaffold structure and measurements of membrane fluidity to determine whether stabilization tracks bulk membrane rigidification. Differential fractionation of plasma and organellar membranes could determine whether the preferential rescue of oIMPs persists in physically resolved membrane systems. But as is, our findings reframe CHS not only as a ligand engaging discrete cholesterol-binding pockets, but as a broadly acting sterol scaffold that stabilizes IMPs in a concentration-dependent and sequence-independent manner, with its magnitude shaped by transmembrane architecture and a pronounced preference for the most thermally vulnerable populations. More broadly, this work establishes MM-TPP as a generalizable platform for resolving lipid-dependent effects on IMP stability at proteome scale and provides a framework for systematically investigating membrane additives and sterol analogs whose mechanisms of action remain largely assumed rather than experimentally resolved.

## Limitations of the study

Several methodological differences warrant consideration. First, CHS-induced stabilization in E. coli membranes was assessed by SDS-PAGE densitometry rather than LC-MS/MS, providing lower proteome coverage and quantitative resolution than the mouse liver datasets. Second, because CHS stocks were prepared in DDM, increasing CHS concentrations also increased the detergent concentration above the nominal 1% target. However, the persistence of CHS-dependent effects in detergent-free Peptidisc and nanodisc assemblies, the lack of similar responses to DOPC or sphingomyelin, and the comparable stabilization observed when CHS was added before or after detergent solubilization argue against this difference being the principal driver. Third, our lipid screen was limited to CHS, DOPC, and sphingomyelin; other lipid species such as phosphatidylethanolamine (PE), phosphatidylserine (PS), and phosphatidylinositol (PI) are known to influence the stability and function of many membrane proteins and were not assessed here. It is therefore possible that these or other lipids could exert additional, protein-specific stabilizing or destabilizing effects at the proteome level that were not captured in the present study. Finally, the DDM and Peptidisc datasets were acquired using different MS acquisition strategies (DDA and DIA, respectively), introducing an analytical confound in addition to the membrane mimetic difference between the systems.

## Acknowledgements

We thank Dr. Huan Bao and Dr. Martin Spiess for the generous gifts of the spNW30 nanodisc plasmid (Addgene #173485) and APEX2 plasmid (Addgene #109420), respectively. Work in the Duong lab was supported by an NSERC Discovery grant. Work in the Babu lab was supported by the Canada Foundation for Innovation and CIHR Foundation grant FDN-154318. A. B. is supported by a UBC 4-year fellowship, an Amplify Doctoral Award from Triangle (Training a new generation of researchers in gastroenterology and liver), and a Mast3 (Mass Spectrometry Team Training and Transition) scholarship.

## Notes

The authors declare the following competing financial interest(s): FDVH is the scientific founder of Peptidisc Biotech. The mass spectrometry proteomics data have been deposited to the ProteomeXchange Consortium via the PRIDE partner repository^48^.

Proteome profiling DIA data (Figure 1): **Project accession:** PXD080996 **Reviewer access details**

Log in to the PRIDE website using the following details:

**Project accession:** PXD080996

**Token:** GvCBQvQO51CH

SP4 DDA data (Figure 4, Table 2, Figure 5) - **Project accession:** PXD080879 **Reviewer access details**

Log in to the PRIDE website using the following details:

**Project accession:** PXD080879

**Token:** u5lhDRZEU9NG

Peptidisc TPP DIA data (Figure 4, Table 2, Figure 5): **Project accession:** PXD081013

**Reviewer access details**

Log in to the PRIDE website using the following details:

**Project accession:** PXD081013

**Token:** w0tHMqvdZQuT

## Author contributions

Conceptualization: A.B., F.A., F.D.v.H.; Formal analysis: A.B.; Supervision: F.D.v.H., M.B.; Funding acquisition: A.B., F.D.v.H., M.B.; Validation: A.B.; Investigation: A.B., S.C., H.A. F.A.; Visualization: A.B., S.C.; Methodology: A.B., F.A.; Writing - original draft: A.B.; Project administration: A.B., F.D.v.H.; Writing - review and editing: A.B., F.D.v.H.

| REAGENT or RESOURCE | SOURCE | IDENTIFIER |
| --- | --- | --- |
| **Bacterial and virus strains** |  |  |
| *E. coli* BL21 (DE3) | Lab stock | N/A |
| **Mammalian tissue** |  |  |
| C57BL/6J male mice | UBC Centre for Comparative Medicine | N/A |
| **Chemicals, peptides, and recombinant proteins** |  |  |
| Peptidisc | Peptidisc Biotech | www.peptidisc.com |
| His-Peptidisc | Peptidisc Biotech | www.peptidisc.com |
| PDET-1 | Peptidisc Biotech | www.peptidisc.com |
| Dodecyl-β-d-maltoside (DDM) | Bioshop | DDM001.5 |
| Phosphate Buffered Saline pH 7.4 | Life Technologies | 10010023 |
| Tris | Bio Shop | TRS003.5 |
| NaCl | Fisher Scientific | BP358-212 |
| Ethylenediaminetetraacetic acid (EDTA) | BioShop | EDT002.500 |
| cOmplete protease inhibitor cocktail | Sigma-Aldrich | 11697498001 |
| Phenylmethylsulfonyl fluoride (PMSF) | Sigma-Aldrich | P-7626 |
| MgCl<sub>2</sub> | Sigma-Aldrich | M8266-1KG |
| DNase I | Life Technologies | 18047019 |
| Glycerol | Fisher Scientific | BP229-4 |
| Kanamycin Monosulfate | GoldBio | K-120-5 |
| IPTG | GoldBio | I2481C5 |
| Ampicillin (Sodium) | GoldBio | A-301-SL10 |
| L-Arabinose | GoldBio | A-300-25 |
| Ni-NTA agarose | Qiagen | 30210 |
| Dithiothreitol (DTT) | BioShop | DTT002.10 |
| Yeast Extract | BioShop | YEX555.500 |
| Tryptone | BioShop | TRP402.5 |
| Imidazole | BioShop | IMD510.500 |
| Acrylamide (40%) | BioShop | ACR006.500 |
| Bis-acrylamide (2%) | BioShop | ACR007.500 |
| TEMED | BioShop | TEM001.25 |
| Ammonium persulfate | Fisher Scientific | BP179-25 |
| Coommassie blue G-250 | GoldBio | C-460-50 |
| 100% Ethanol | Fisher Scientific | A962<sup>F</sup>-1Gal |
| Glacial Acetic Acid | Fisher Scientific | 3512694 |
| Urea | BioShop | URE002.5 |
| Iodoacetamide (IAA) | BioShop | IOD500.5 |
| 100K MWCO filter units | Millipore-Sigma | UFC910024 |
| MS grade proteases | Thermo Fisher Scientific | 90057 |
| C18-HD disk 47 mm | Empore | 41120000 |
| Polygoprep 300-20 C18 (C18 resin) | Macherey-Nagel | 711025.100 |
| Methanol | Sigma-Aldrich | 179337-4L |
| Glass Spheres | Sigma-Aldrich | 440345-100G |
| Acetonitrile | Fisher Scientific | A211 |
| Sodium Dodecyl Sulfate | Sigma-Aldrich | 75746-250G |
| Water suitable for HPLC | Sigma-Aldrich | 270733-1L |
| Hydrochloric acid (HCl) solution | Fisher Scientific | A144-212 |
| Sodium hydroxide (NaOH) | Fisher Scientific | S318-3 |
| ACRYLAMIDE/BIS-<br>ACRYLAMIDE, 40% Solution<br>37.5:1 | Bioshop | ACR005.500 |
| Ammonium bicarbonate | Fisher Scientific | A643500 |
| Formic acid | Sigma-Aldrich | 695076-100ML |
| Brain Sphingomyelin | Avanti Research | 860062P-25mg |
| 1 2-dioleoyl-sn-glycero-3-phosphocholine (DOPC) | Avanti Research | 850375C |
| Cholesteryl hemisuccinate | Avanti Research | 850524P-1g |
| Bovine Serum Albumin | Fisher Scientific | BP671-1 |
| Bradford Assay Solution | BioRad | 5000006 |
| Chloroform | Fisher Scientific | C2984 |
| **Plasmids** |  |  |
| pET28-his<sub>6</sub>MsbA | Lab stock | N/A |
| pBAD22-his<sub>6</sub>YidC | Lab stock | N/A |
| pBAD33-his<sub>6</sub>YibN | Lab stock | N/A |
| pBAD33-his<sub>6</sub>YibN-TMS-APEX2 | Generated in this study | N/A |
| pET24a-VHH-APEX2 | Addgene | 109420 |
| pET28a-his<sub>6</sub>spNW30 | Addgene | 173485 |

| Software and algorithms |  |  |
| --- | --- | --- |
| ImageJ (v1.54t) | ImageJ | www.imagej.net |
| Rstudio (v2026.5.0.218) | Posit | https://posit.co/ |
| Fragpipe (v24.0) | Nesvi Lab | https://fragpipe.nesvilab.org/ |
| DIA-NN (v2.0) | Vadim Demichev | https://github.com/vdemichev/diann |
| Phobius | Stockholm Bioinformatics Centre | https://phobius.sbc.su.se/ |
| Biorender | Biorender | https://www.biorender.com/ |
| Uniprot | UniProt Consortium | https://www.uniprot.org/ |

## Methods

### Bacterial Strains and Plasmids

*E. coli* strain BL21 (DE3) was from our laboratory collection. Plasmids encoding His-tagged MsbA (pET28-his_6_MsbA)<u>^22^</u>, His-tagged YidC (pBAD22-his6YidC)<u>^49^</u>, and His-tagged YibN (pBAD33-his_6_YibN)<u>^49^</u> were previously constructed in our laboratory. The pET24a-VHH-APEX2 plasmid was a gift from Dr. Martin Spiess (Addgene plasmid #109420). The pET28a-his_6_spNW30 plasmid was a gift from Dr. Huan Bao (addgene plasmid #173485). The pBAD33-his_6_YibN-TMS-APEX2 plasmid was generated in this study by PIPE cloning<u>^50^</u>, joining the YibN transmembrane segment (N_ter_-MQEIMQFVGRHPILSIAWIALLVAVLVTT-Cter), the linker sequence (N_ter_-FKSLTSKVKVITRGEATRLINKEDAVVVDLRQRDDFRKGH-Cter), and the APEX2 sequence. The construct was confirmed by Sanger sequencing (Genewiz). Primers used for cloning are listed in Supplemental Table 4.

### Mouse Liver Membrane Fraction Preparation

Animal studies were approved by the University of British Columbia with assurance from Animal Care Services, Centre for Comparative Medicine. Eight-month-old C57BL/6J male mice were euthanized by CO_2_ asphyxiation and livers were immediately dissected. The excised livers were washed three times in ice-cold PBS to remove residual blood, then minced and homogenized in ice-cold hypotonic lysis buffer (10 mM Tris–HCl pH 7.4, 30 mM NaCl, 1 mM EDTA, 1× cOmplete protease inhibitor cocktail and 1 mM PMSF) using a tight-fitting Dounce homogenizer. All subsequent steps were performed at 4°C or on ice. MgCl_2_ and DNase I were added to final concentrations of 10 mM and 50 µg/mL, respectively, and the suspension was further homogenized and incubated for 10 min. The lysate was passed through a French press three times at 500 psi at 4°C. Unbroken cells and the nuclear fraction were removed by low-speed centrifugation (1,200g, 10 min, 4°C), and the crude membrane fraction was collected by ultracentrifugation (135,520g, 45 min, 4°C) in a Beckman TLA110 rotor. The membrane pellet was resuspended in 100 µL of TSG buffer (50 mM Tris pH 7.4, 100 mM NaCl, 10% glycerol) and stored at −70°C until use.

### Protein Expression and Membrane Fraction Preparation

Proteins were expressed in *E. coli* BL21 (DE3) grown in LB medium. For MsbA expression, cultures were grown in LB supplemented with kanamycin (50 µg/mL) at 37°C to an OD600 of ∼0.4, upon which expression was induced with 0.5 mM IPTG for 4 h at 37°C. For YidC and YibN-TMS-APEX2 expression, cultures were grown in LB supplemented with ampicillin (100 µg/mL) at 37°C to an OD600 of ∼0.4, and expression was induced with 0.2% (w/v) arabinose for 4 h at 37°C. For spNW30, a soluble nanodisc scaffold protein, cultures were grown in LB supplemented with kanamycin (50 µg/mL) at 37°C to an OD600 of ∼0.4, and expression was induced with 0.2 mM IPTG overnight at 16°C. Following expression, cells were harvested by centrifugation (6,000g, 6 min, 4°C) and resuspended in TSG supplemented with 1 mM PMSF. Cells were lysed by three passes through a microfluidizer at 15,000 psi at 4°C. Unbroken cells and large aggregates were removed by low-speed centrifugation (4,350g, 10 min, 4°C), and the crude membrane fraction was isolated by ultracentrifugation (125,440g, 45 min, 4°C). For MsbA, YidC, and YibN-TMS-APEX2, the crude membrane pellet was resuspended in TSG at 5 mg/mL and stored at −20°C until use. For spNW30, the soluble supernatant fraction was subsequently Ni-NTA purified.

### Lipid Preparation

Sphingomyelin, DOPC, and CHS were used as supplied by the manufacturer. Each lipid was dissolved in chloroform at 10 mg/mL and dried to a thin film under a gentle stream of nitrogen. Dried lipid films were resuspended in TS (50 mM Tris pH 7.4 and 100 mM NaCl) containing 0.5% (w/v) DDM to a final concentration of 5 mg/mL by vortexing for 5 min followed by water bath sonication for 15 min at room temperature. Stocks were aliquoted and stored at −70°C until use.

### Lipid Supplementation of Membrane Proteome Library

Lipid supplementation was performed either prior to or concurrent with DDM solubilization of crude membranes. Concurrent supplementation was used for all proteomics and TPP experiments; the pre-solubilization approach was evaluated as an experimental control. For concurrent supplementation, the indicated lipid (500 µg for the initial membrane proteome profiling experiment; 0.5–4 mg CHS for TPP experiments) was added together with 1% (w/v) DDM to 1 mg of crude membrane and incubated for 45 min at 4°C with gentle agitation. Insoluble material was removed by ultracentrifugation (135,520g, 15 min, 4°C) and the clarified extract was collected as the supernatant. Note that because CHS stocks were prepared in 0.5% (w/v) DDM, increasing CHS concentrations introduced proportionally higher DDM above the nominal 1% target. For pre-solubilization supplementation, 1 mg of crude membrane was incubated with the indicated lipid for 45 min at 4°C with gentle agitation. Membranes were pelleted by ultracentrifugation (135,520g, 15 min, 4°C) to remove unincorporated lipids, and the supernatant was discarded. The membrane pellet was resuspended and solubilized in 1% (w/v) DDM for 45 min at 4°C to extract membrane proteins, followed by clarification at 135,520g for 15 min at 4°C to collect the solubilized extract.

### Reconstitution of Membrane Proteome into Peptidisc and Nanodiscs

Two reconstitution protocols were used depending on the experimental context: an “on-filter” approach for proteome-level library preparation and an “on-bead” approach for single-protein reconstitution from *E. coli* crude membranes. For on-filter reconstitution, crude membranes (∼1 mg) were solubilized in an ice-cold TS buffer supplemented with 1% (w/v) DDM for 45 min at 4°C with gentle shaking. For lipid-supplemented conditions, the DDM extract was prepared as described in the Lipid Supplementation section. The detergent extract was clarified by ultracentrifugation (135,520g, 15 min, 4°C), and 1 mg aliquots were incubated with a 2-fold molar excess of His_6_-tagged Peptidisc or a 1-fold molar excess of His_6_-tagged spNW30, calculated relative to the total protein concentration of the DDM extract as determined by Bradford assay, for 15 min at RT. Insoluble material was removed by ultracentrifugation (135,520g, 15 min, 4°C), and the reconstituted sample was transferred to a 100 kDa MWCO centrifugal filter unit and diluted to 5 mL in TS buffer. To reduce DDM to sub-micellar concentrations and drive Peptidisc assembly, the sample was concentrated at 3,000g for 20 min to approximately 200 µL, diluted again to 5 mL, and concentrated a second time to approximately 200 µL. For proteome profiling experiments, the reconstituted library was subsequently affinity purified by incubation with 100 µL of Ni-NTA resin for 1 h at 4°C with shaking, followed by five washes with 1 mL TS buffer containing 20 mM imidazole, and elution in 150 µL of TSG buffer containing 600 mM imidazole. For MM-TPP experiments, the Ni-NTA purification step was omitted; following DDM dilution, the reconstituted sample was used directly to preserve the complexity of the membrane proteome and avoid any affinity-based selection that could bias thermal stability measurements. For on-bead reconstitution, all steps were carried out at 4°C and untagged Peptidisc scaffold was used in place of His_6_-tagged Peptidisc, as affinity purification in this protocol is directed through the histidine tag on the target protein rather than the scaffold. Crude membranes (2 mg) were solubilized in TS buffer supplemented with 1% (w/v) DDM and CHS for 45 min, and insoluble aggregates were removed by ultracentrifugation (135,520g, 15 min, 4°C). For *E. coli* MM-TPP experiments, 4 mg CHS was added during solubilization; for the PDET-1 experiment, 1 mg CHS was used. The clarified detergent extract was incubated with pre-equilibrated Ni-NTA resin (150 µL) for 1 h on a tabletop rocker. The resin was pelleted by centrifugation (2000g, 2 min, 4°C) and washed twice with TS buffer containing 20 mM imidazole and 0.02% (w/v) DDM, then resuspended in 1 mL TS buffer supplemented with 1 mg untagged Peptidisc scaffold and washed five times with TS buffer. For the PDET-1 experiment, 1 mg PDET-1 was used in place of untagged Peptidisc scaffold for reconstitution; all other steps were identical. Peptidisc-reconstituted proteins were eluted in 150 µL TSG buffer containing 600 mM imidazole and stored at −20°C until use. For both protocols, all fractions from reconstitution through elution were analyzed by SDS-PAGE with Coomassie blue staining to confirm reconstitution efficiency and purity.

### Membrane Mimetic-Thermal Proteome Profiling Assay

Membrane protein libraries reconstituted in Peptidisc or solubilized in DDM, with or without CHS, were quantified by Bradford assay. For each sample, 150 µg of protein was transferred to a PCR tube and brought to a final volume of 50 µL with a TS buffer. A RT reference sample was prepared identically and carried through all downstream steps without heat treatment. Samples were subjected to thermal denaturation in a thermocycler for 3 minutes at each temperature point, with the heated lid set to the maximum temperature of the gradient for each experiment. For proteome-level experiments, temperature gradients spanning RT to 90°C (DDM conditions) or RT to 100°C (Peptidisc conditions) were applied, with specific temperature points for each experiment as indicated in the figure legends. For single-protein experiments, MsbA was profiled over a lower temperature range (30–55°C) to match its lower thermal stability, and YibN-TMS-APEX2 reconstituted in Peptidisc was subjected to isothermal incubation at 90°C for durations of 0, 3, 5, 10, 15, and 20 minutes rather than a temperature gradient. Following heat treatment, aggregated proteins were pelleted by ultracentrifugation (135,520g, 15 min, 4°C) and the clarified supernatant was collected. Proteins were resolved by SDS-PAGE. For MS-based experiments, DDM-solubilized samples underwent SP4-based protein capture and Peptidisc-reconstituted samples were processed directly, as described in the Mass Spectrometry Sample Preparation section.

### Densitometry

All densitometry was performed in ImageJ (v 1.54t) on raw .gel image files. A fixed rectangular ROI was sized to encompass the largest band in each gel series and applied uniformly across all lanes without adjustment. Where gel images were captured with dark background polarity, pixel values were inverted prior to measurement. Mean gray values were recorded, and background was subtracted using a constant value measured from a protein-free region of the same gel. All values within a series were normalized to the corresponding RT lane value, set to 1. Data were visualized using ggplot2 in RStudio (v2026.5.0.218).

### Native Gel Electrophoresis

Equal volumes of 4% and 12% acrylamide solutions were prepared in advance, and linear gradient gels were poured manually. The cross-linking agents, TEMED and ammonium persulfate, were added immediately before gradient mixing. Plastic combs (Bio-Rad) were then inserted, and the gels were allowed to polymerize for 30 min before storage at 4 °C. For clear native gel electrophoresis, the anode and cathode buffers consisted of buffer N (37 mM Tris–HCl pH 8.8, 35 mM Glycine). For blue-native gel electrophoresis, the anode buffer was replaced with buffer NA (Buffer N, 180 μM Coomassie Blue G-250), while the cathode buffer remained unchanged^51^.

### Mass spectrometry Sample Preparation

Two sample preparation workflows were used depending on the solubilization platform. DDM-solubilized samples underwent SP4-based protein capture prior to digestion; Peptidisc-reconstituted samples were processed directly. For SP4-based preparation, glass spheres (9–13 µm diameter) were resuspended in water, washed once with 100% acetonitrile, rinsed twice with water, and resuspended at 50 mg/mL; beads were isolated by centrifugation at 16,000g for 1 min after each wash. Protein aliquots of 100 µg were gently vortex-mixed with 1 mg silica beads, acetonitrile was added to a final concentration of 80%, and samples were centrifuged at 16,000g for 5 min. Beads were rinsed three times with 500 µL of 80% ethanol without disturbing the pellet, then resuspended in 100 µL of 6 M urea. Beads were removed by centrifugation at 16,000g for 1 min and the protein-containing supernatant in 6 M urea was carried forward to the shared digestion protocol described below, beginning at the DTT reduction step. For Peptidisc-reconstituted samples, 100 µg of protein was denatured in 6 M urea at RT for 30 min before proceeding to digestion. All samples were reduced with 10 mM DTT for 1 h at RT, alkylated with 20 mM IAA in the dark at RT for 30 min, and quenched with 10 mM DTT for 30 min at RT. Urea was diluted to 1 M with 50 mM ammonium bicarbonate (pH 8.0), and MS-grade trypsin (Thermo Fisher Scientific, cat. 90057) was added at a 1:100 enzyme-to-protein ratio for 24 h at 25°C with shaking. Digested peptides were acidified with 10% (v/v) formic acid to pH 3, desalted using hand-packed C18 Stage-Tips (Empore C18 material), eluted with 40% acetonitrile in 0.1% formic acid, and dried by vacuum centrifugation.

### Liquid Chromatography and Mass Spectrometry Analysis

NanoLC connected to an Orbitrap Exploris 240 mass spectrometer (Thermo Fisher Scientific) was used for the analysis of all samples. The peptide separation was carried out using a Proxeon EASY nLC 1200 System (Thermo Fisher Scientific) fitted with a custom-made C18 column (15 cm x 150 μm ID) packed with HxSil C18 3 μm Resin 100 Å (Hamilton). A gradient of water/acetonitrile/0.1% formic acid was employed for chromatography. The samples were injected onto the column and run for 180 minutes at a flow rate of 0.60 μl/min. The peptide separation began with 1% acetonitrile, increasing to 3% in the first 4 minutes, followed by a linear gradient from 3% to 23% acetonitrile over 86 minutes, then another increase from 24% to 80% acetonitrile over 35 minutes, and finally a 35-minute wash at 80% acetonitrile, and then decreasing to 1% acetonitrile for 10 min and kept 1% acetonitrile for another 10 min. The eluted peptides were ionized using positive nanoelectrospray ionization (NSI) and directly introduced into the mass spectrometer with an ion source temperature set at 250°C and an ion spray voltage of 2.1 kV. In the case of data-dependent acquisition (DDA) mode, full-scan MS spectra (m/z 350–2000) were captured in Orbitrap Exploris 240 at a resolution of 120,000 (m/z 400). The automatic gain control was set to 1e6 for full FTMS scans and 5e4 for MS/MS scans. Ions with intensities above 1500 counts underwent fragmentation via NSI in the linear ion trap. The top 15 most intense ions with charge states of ≥2 were sequentially isolated and fragmented using normalized collision energy of 30%, activation Q of 0.250, and an activation time of 10 ms. Ions selected for MS/MS were excluded from further selection for 3 seconds. For EASY-nLC DIA, Orbitrap full MS scans were acquired from 375 to 1500 m/z at a resolution of 120,000 at m/z 400 with a normalized automated gain control (AGC) target of 300% and a maximum ion injection time set to Auto. For MS/MS scans, the HCD collision energy was set to 30%, the Orbitrap resolution to 15,000 at m/z 400, the normalized AGC target to 800%, the maximum injection time set to Auto, and the mass range to m/z 375 to 1500. For a theoretical cycle time of 3 s, 70 DIA windows of 19 m/z and an overlap of 0.5 m/z were used. The desired minimum points across the peak were set to 9, and the data type was centroid. The Orbitrap Exploris 240 mass spectrometer was operated using Thermo XCalibur software.

### Mass Spectrometry Data Analysis

Raw MS data were processed using two pipelines depending on acquisition mode. DDA data from DDM-solubilized samples were analyzed in FragPipe; DIA data from Peptidisc-reconstituted samples were analyzed in DIA-NN. Both pipelines used fully tryptic cleavage specificity (Trypsin/P) with a maximum of two missed cleavages, fixed cysteine carbamidomethylation, and a 1% FDR threshold. DDA data were processed using MSFragger (v4.1) within FragPipe (v22.0)^52,53^ under the LFQ-MBR workflow. Database searches were performed against the UniProt *Mus musculus* proteome (UP000000589, December 2024, 54,727 entries) or the UniProt *E. coli* proteome (UP000000625, May 2026, 4,403 entries), each concatenated with its corresponding reverse decoy sequences. Precursor and fragment mass tolerances were set to 20 ppm. Peptide length was constrained to 7–50 amino acids with a mass range of 500–5,000 Da. Variable modifications included methionine oxidation and N-terminal acetylation, with a maximum of three variable modifications per peptide. Peptide-spectrum match and protein-level FDR were each controlled at 1% using MSBooster with Percolator and sequential decoy filtering. Proteins were quantified using MaxLFQ intensities via IonQuant with match-between-runs enabled at 1% ion-level FDR, 10 ppm precursor mass tolerance, and 0.4 min retention time tolerance, using the "unique plus razor" peptide-protein assignment strategy. Intensities were normalized across runs by MaxLFQ. DIA data were processed using DIA-NN (v1.9.1)^54^ in library-free mode with common contaminant sequences included. Spectral libraries were generated from FASTA digests of the *Mus musculus* reference proteome (UP000000589) using deep learning–based prediction of spectra and retention times. No variable modifications were included; N-terminal methionine excision was enabled as a fixed processing step. Peptide length was constrained to 7–30 amino acids with a precursor charge range of 1–4, precursor m/z range of 300–1,800, and fragment ion m/z range of 200–1,800. Protein inference was performed at the gene level using heuristic inference with shared spectra excluded. FDR was controlled at 1% at the precursor level. Quantification used the QuantUMS high-precision strategy with RT-dependent cross-run normalization. The neural network classifier was run in single-pass mode, and all datasets were processed within a single DIA-NN run to ensure consistent quantification and normalization across samples.

### Protein Annotation

The protein list obtained from DIA-NN and Fragpipe was subjected to a gene ontology (GO)-term analysis using the UniProtKB database^55^ to identify proteins with the GO-term “membrane”. Within this group, proteins containing at least one α-helical transmembrane segment were labeled as IMPs. While those without any transmembrane segment were annotated as MAPs. Proteins not annotated with the "membrane" GO term were classified as soluble. The Phobius web server was utilized (http://phobius.sbc.su.se/) to predict the number of transmembrane segments^56^. The subcellular localization of the IMPs was further classified using the GO-term ‘Subcellular location [CC].’ These were divided into pIMPs if they contain the GO-term ‘plasma membrane’. Alternatively, they were labeled as oIMPs when containing keywords such as ‘ER, Golgi membrane, vesicle membrane (including endosome, exosome, peroxisome, lysosome, and vesicle), mitochondrial, and nucleus membranes’.

### Figure Generation, Data Analysis, and Code Availability

The graphical abstract was created with BioRender.com. Figures 1, 2B, 3B, 4-5, 6B, 7B, Table 1-2, Supplemental Table 1-3, Supplemental Figures 2-5, 6B, 13B, 14B were generated in R (v4.5.1) using RStudio (v2026.5.0.218) with the following packages: dplyr, tidyverse, rstatix, tidyr, stringr, multcomp, httr, jsonlite, ggplot2, ggrepel, ggsignif, and ggh4x. All data analysis was performed in RStudio. Custom scripts used for data processing and visualization are available from the lead contact upon reasonable request.

---

**Supplemental Figure 1.**
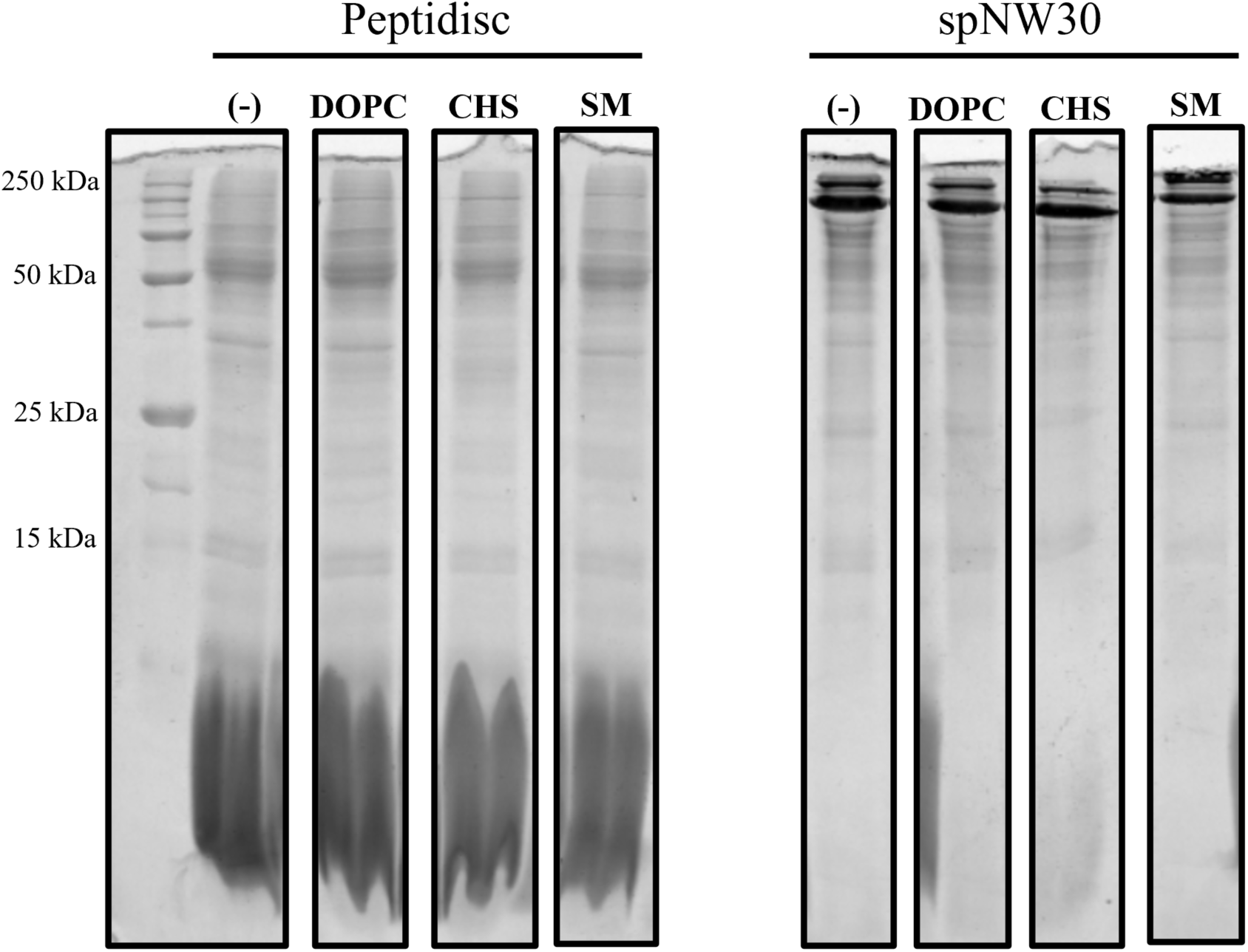
SDS-PAGE of mouse liver membrane proteome libraries reconstituted into Peptidisc or spNW30 nanodiscs with lipid additives. Mouse liver crude membranes were solubilized in DDM supplemented with 500 µg of dioleoylphosphatidylcholine (DOPC), cholesteryl hemisuccinate (CHS), or sphingomyelin (SM), followed by reconstitution in Peptidisc (left) or spNW30 nanodiscs (right). Equal volumes of each reconstituted library were resolved by SDS-PAGE and stained with Coomassie. Molecular weight markers are indicated on the left (kDa).

**Supplemental Table 1.** Protein identification counts by subcellular localization across the mouse liver Peptidisc thermal proteome profiling experiment, in the presence or absence of 2 mg CHS. Mouse liver membrane proteins were solubilized in DDM in the presence or absence of 2 mg CHS, reconstituted into Peptidisc, and subjected to thermal proteome profiling across the TPP gradient (RT, 50, 60, 70, 80, 90, 100°C), with samples analyzed by DIA LC-MS/MS and processed using DIA-NN. For each temperature and condition, the total number of detected protein groups (TP) is reported alongside the count and percentage of proteins assigned to each localization category: soluble proteins (SP), membrane-associated proteins (MAP), total integral membrane proteins (tIMP, comprising oIMP and pIMP), organellar integral membrane proteins (oIMP), and plasma membrane integral membrane proteins (pIMP). TP reflects the number of proteins with a detectable intensity value at that specific temperature within each condition; percentages are calculated relative to the total detected protein count for that temperature. Protein localization assignments and tabulation were performed using dplyr and rendered with knitr::kable. CHS, cholesteryl hemisuccinate.

|  | TP | TPs |  |  | tIMPs |  |
| --- | --- | --- | --- | --- | --- | --- |
|  |  | SP | MAP | tIMP | oIMP | pIMP |
| RT | 1613 | 771 (48%) | 426 (26%) | 416 (26%) | 239 (15%) | 177 (11%) |
| 50 | 1415 | 674 (48%) | 368 (26%) | 373 (26%) | 196 (14%) | 177 (12%) |
| 60 | 1068 | 503 (47%) | 276 (26%) | 289 (27%) | 135 (13%) | 154 (14%) |
| 70 | 880 | 405 (46%) | 226 (26%) | 249 (28%) | 107 (12%) | 142 (16%) |
| 80 | 796 | 361 (45%) | 203 (26%) | 232 (29%) | 96 (12%) | 136 (17%) |
| 90 | 690 | 317 (46%) | 179 (26%) | 194 (28%) | 80 (12%) | 114 (16%) |
| 100 | 668 | 315 (47%) | 174 (26%) | 179 (27%) | 70 (11%) | 109 (16%) |
| 2 mg CHS<br>RT | 1775 | 805 (45%) | 475 (27%) | 495 (28%) | 306 (17%) | 189 (11%) |
| 2 mg CHS<br>50 | 1652 | 746 (45%) | 448 (27%) | 458 (28%) | 281 (17%) | 177 (11%) |
| 2 mg CHS<br>60 | 1677 | 753 (45%) | 448 (27%) | 476 (28%) | 284 (17%) | 192 (11%) |
| 2 mg CHS<br>70 | 1540 | 678 (45%) | 408 (26%) | 454 (29%) | 252 (16%) | 202 (13%) |
| 2 mg CHS<br>80 | 1403 | 607 (43%) | 381 (27%) | 415 (30%) | 230 (17%) | 185 (13%) |
| 2 mg CHS<br>90 | 1191 | 516 (44%) | 327 (27%) | 348 (29%) | 182 (15%) | 166 (14%) |
| 2 mg CHS<br>100 | 1140 | 498 (43%) | 316 (28%) | 326 (29%) | 166 (15%) | 160 (14%) |

**Supplemental Figure 2.**
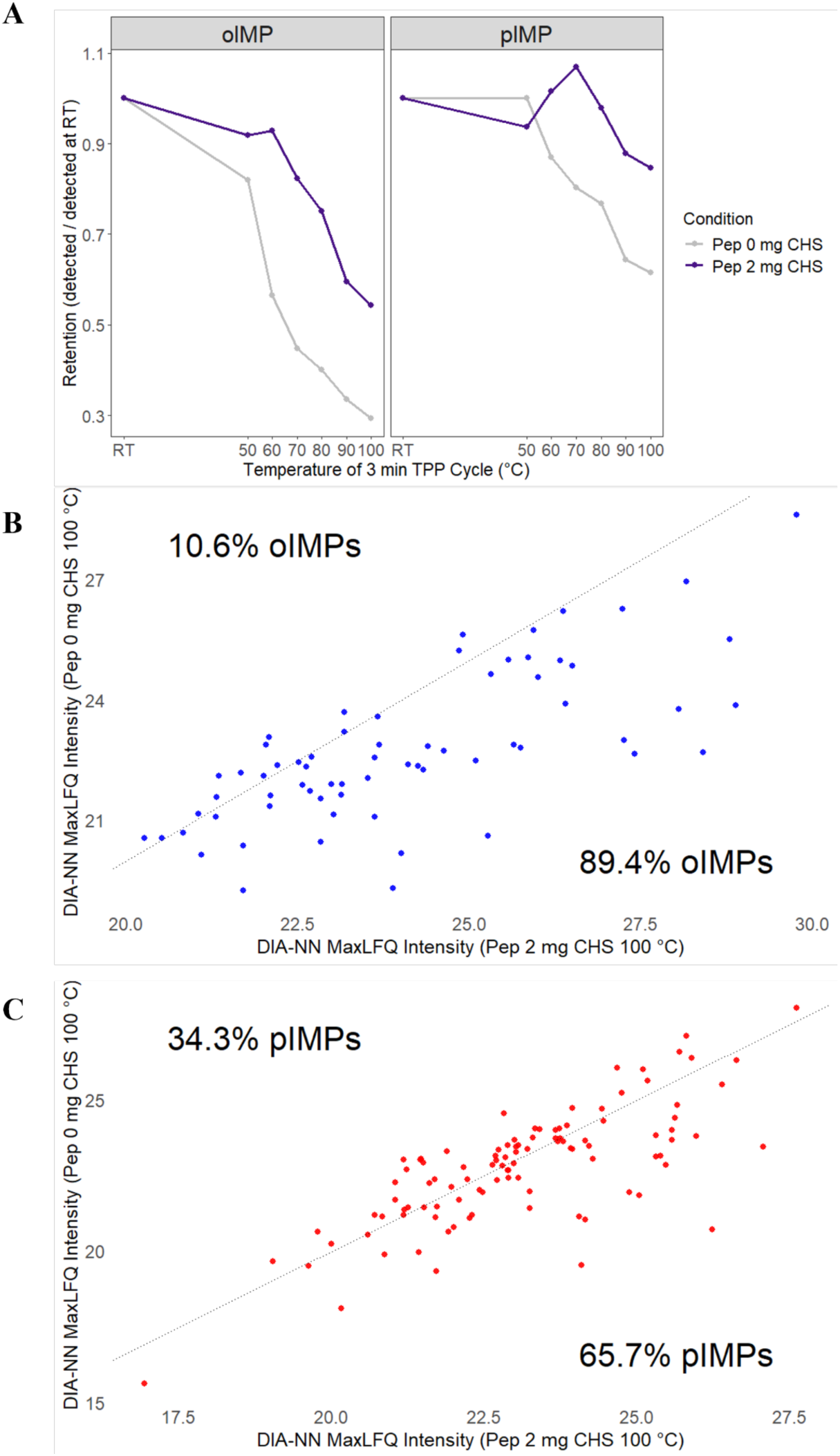
Effect of CHS Supplementation on Thermal Profiles of Organellar and Plasma Membrane Integral Proteins in the Mouse Liver Peptidisc Proteome. The mouse liver membrane proteome was reconstituted in Peptidisc in the presence (2 mg CHS) or absence (0 mg CHS) of CHS and subjected to thermal proteome profiling. (**A**) Proteome-level retention curves for organellar integral membrane proteins (oIMP) and plasma membrane integral membrane proteins (pIMP). For each protein with a valid log2-transformed DIA-NN MaxLFQ intensity at RT, intensity values at each temperature were normalized to the RT value to generate a per-protein relative retention. These were then averaged across all proteins within each subcategory (oIMP or pIMP) to produce a mean retention curve per condition. Curves were computed using dplyr and visualized with ggplot2. (**B–C**) Protein-level intensity scatter plots comparing log2-transformed signal intensities with or without CHS (2 mg) after 3 min incubation at 100°C for oIMPs (B) and pIMPs (C). Points above the diagonal indicate proteins with higher intensity in the 0 mg CHS condition; points below the diagonal indicate proteins with higher intensity in the 2 mg CHS condition). The number and percentage of proteins in each direction are shown relative to all proteins detected in either condition. Pearson correlation coefficients (r) are indicated. Plots were generated using ggplot2 via make_corr_plot_one_group.

**Supplemental Table 2.**
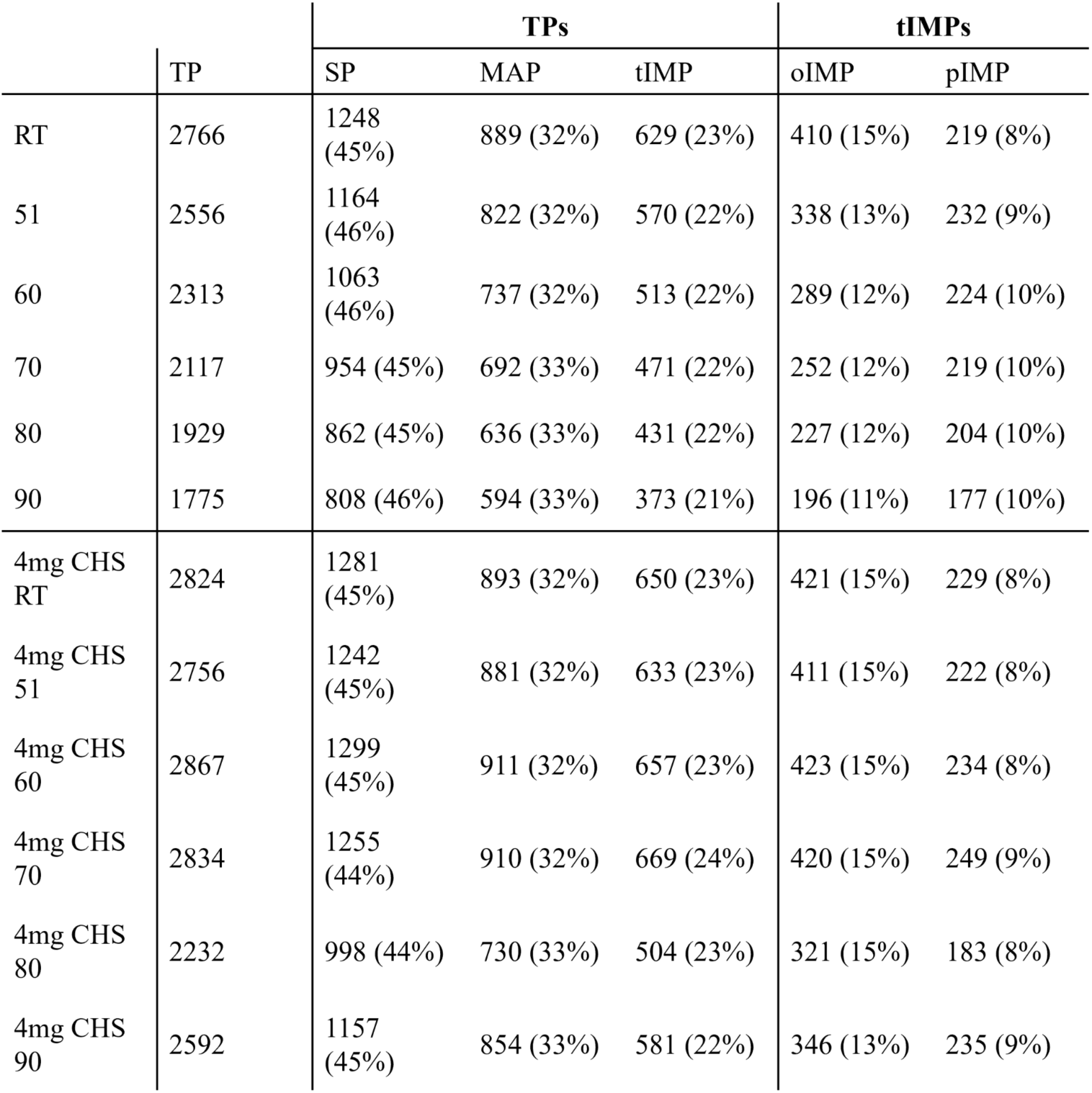
Protein identification counts by subcellular localization across the mouse liver DDM thermal proteome profiling experiment, in the presence or absence of 4 mg CHS. Mouse liver membrane proteins were solubilized in DDM in the presence or absence of 4 mg CHS and subjected to thermal proteome profiling across the TPP gradient (RT, 51, 60, 70, 80, 90°C), prior to SP4 clean-up and DDA LC-MS/MS analysis processed using FragPipe. For each temperature and condition, the total number of detected protein groups (TP) is reported alongside the count and percentage of proteins assigned to each localization category: soluble proteins (SP), membrane-associated proteins (MAP), total integral membrane proteins (tIMP, comprising oIMP and pIMP), organellar integral membrane proteins (oIMP), and plasma membrane integral membrane proteins (pIMP). TP reflects the number of proteins with a detectable intensity value at that specific temperature within each condition; percentages are calculated relative to the total detected protein count for that temperature. Protein localization assignments and tabulation were performed using dplyr and rendered with knitr::kable. CHS, cholesteryl hemisuccinate.

**Supplemental Figure 3.**
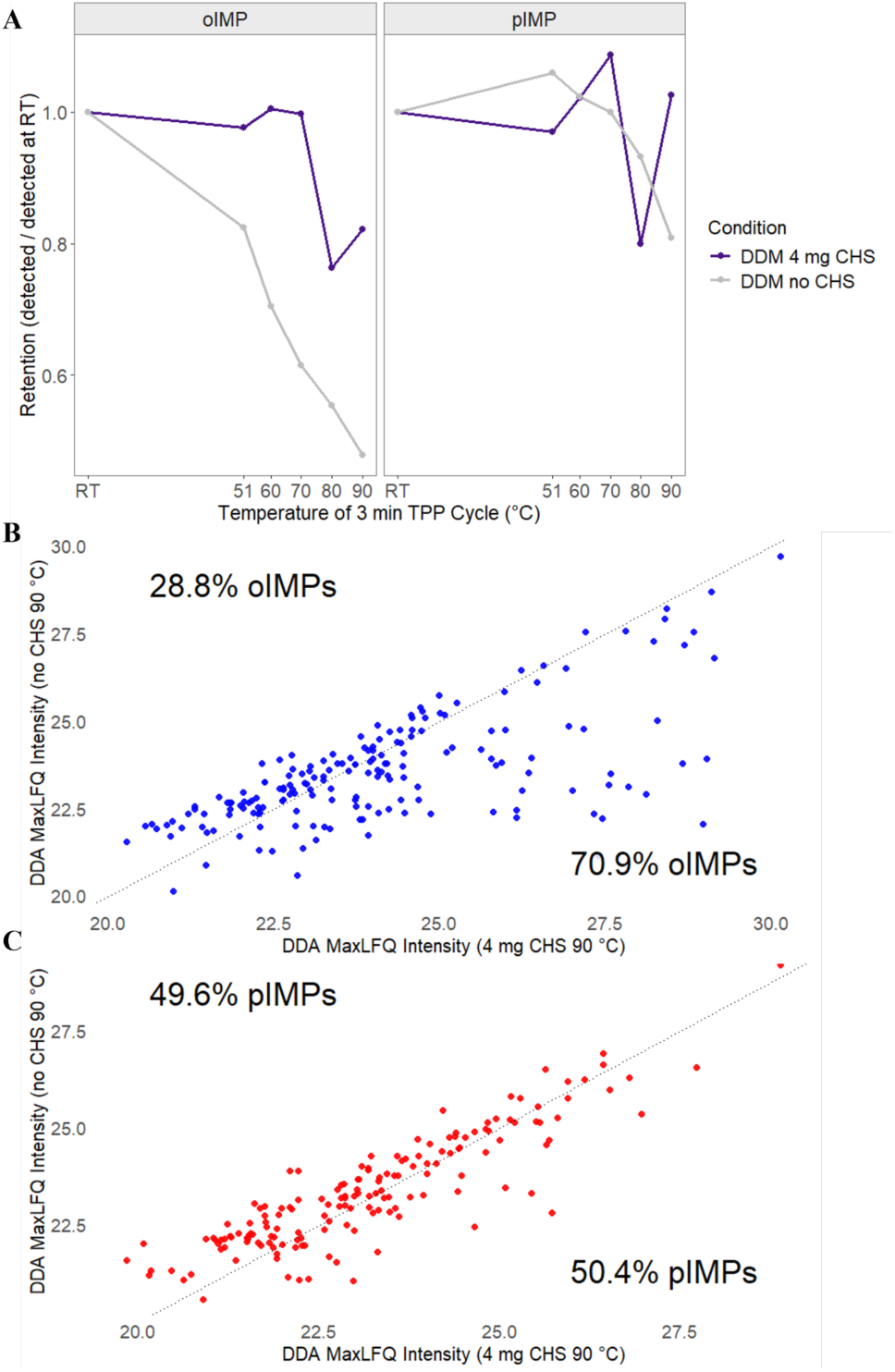
Effect of CHS Supplementation on Thermal Profiles of Organellar and Plasma Membrane Integral Proteins in the Mouse Liver DDM Proteome. Mouse liver membrane proteins were solubilized in DDM with or without 4 mg CHS and subjected to TPP via 3-minute heat treatment at the indicated temperatures. Aggregated proteins were removed by ultracentrifugation, and the soluble supernatant was analyzed by DDA LC-MS/MS. Protein abundance was quantified using log2 transformed MaxLFQ intensity as implemented in FragPipe. (**A**) Proteome-level retention curves for organellar integral membrane proteins (oIMP) and plasma membrane integral membrane proteins (pIMP). For each protein detected at RT, MaxLFQ intensity at each temperature was normalized to its RT value, and the resulting per-protein relative retention was averaged across all proteins within each subcategory to generate a mean retention curve per condition. (**B**) Individual thermal retention curves for the ten transmembrane-containing proteins showing the greatest CHS-dependent stabilization. For each protein, intensity at each temperature was normalized to its RT value and plotted as relative retention. Proteins are ranked from highest to lowest stabilization score, defined as the mean difference in log2 intensity change from RT between the CHS and no CHS conditions across a minimum of three stress temperatures. Plot generated using ggplot2.

**Supplemental Figure 4.**
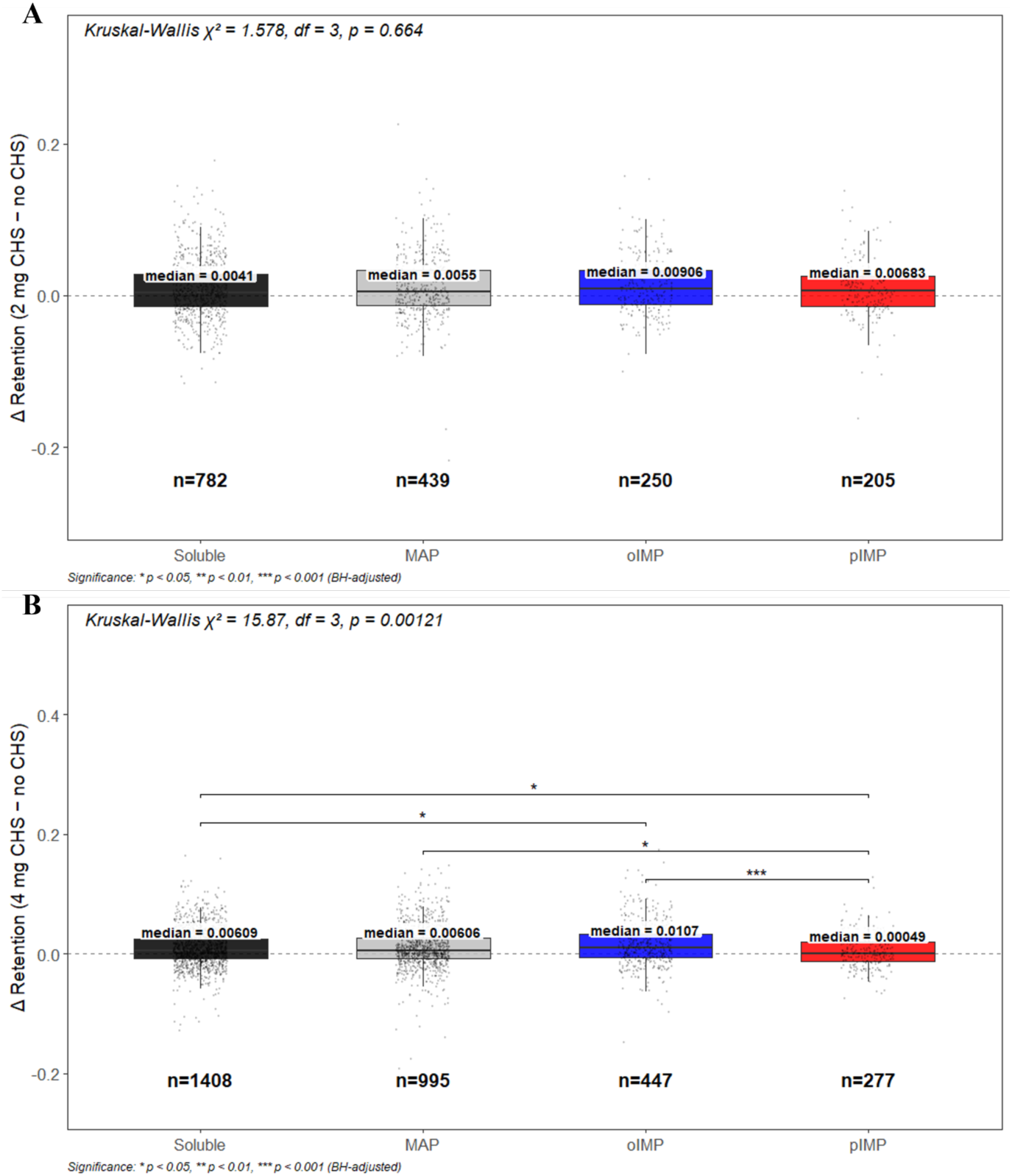
CHS-induced retention shifts differ by membrane protein localization class in DDM but not in Peptidisc. Per-protein retention shift (Δ retention = CHS − no CHS) was compared across four localization classes — Soluble, MAP, oIMP, and pIMP — for DDM and Peptidisc CHS mouse liver LC-MS/MS data. **(A)** In Peptidisc (2 mg CHS − no CHS), Kruskal-Wallis testing showed no significant difference in CHS-induced retention shift across localization classes (χ² = 1.578, df = 3, p = 0.664). Boxplots show median (labeled) and interquartile range; points represent individual proteins; n denotes protein number per class. **(B)** In DDM (4 mg CHS − no CHS), retention shift differed significantly by localization class overall (Kruskal-Wallis χ² = 15.87, df = 3, p = 0.00121). Pairwise BH-adjusted Wilcoxon contrasts showed significant differences between Soluble and oIMP (p = 0.0204), Soluble and pIMP (p = 0.0180), pIMP and MAP (p = 0.0160), and pIMP and oIMP (p = 0.000662), with the oIMP–pIMP contrast retaining significance after correction at the highest stringency; oIMP and MAP (p = 0.0593) and Soluble and MAP (p = 0.637) were not significant. oIMP showed the largest median retention shift (0.0107), while pIMP showed the smallest (0.00049). Statistical analysis was performed in R using kruskal.test and pairwise.wilcox.test (Benjamini-Hochberg adjusted), with data processing in dplyr/tidyr and figures generated using ggplot2 and ggsignif.

**Supplemental Figure 5.**
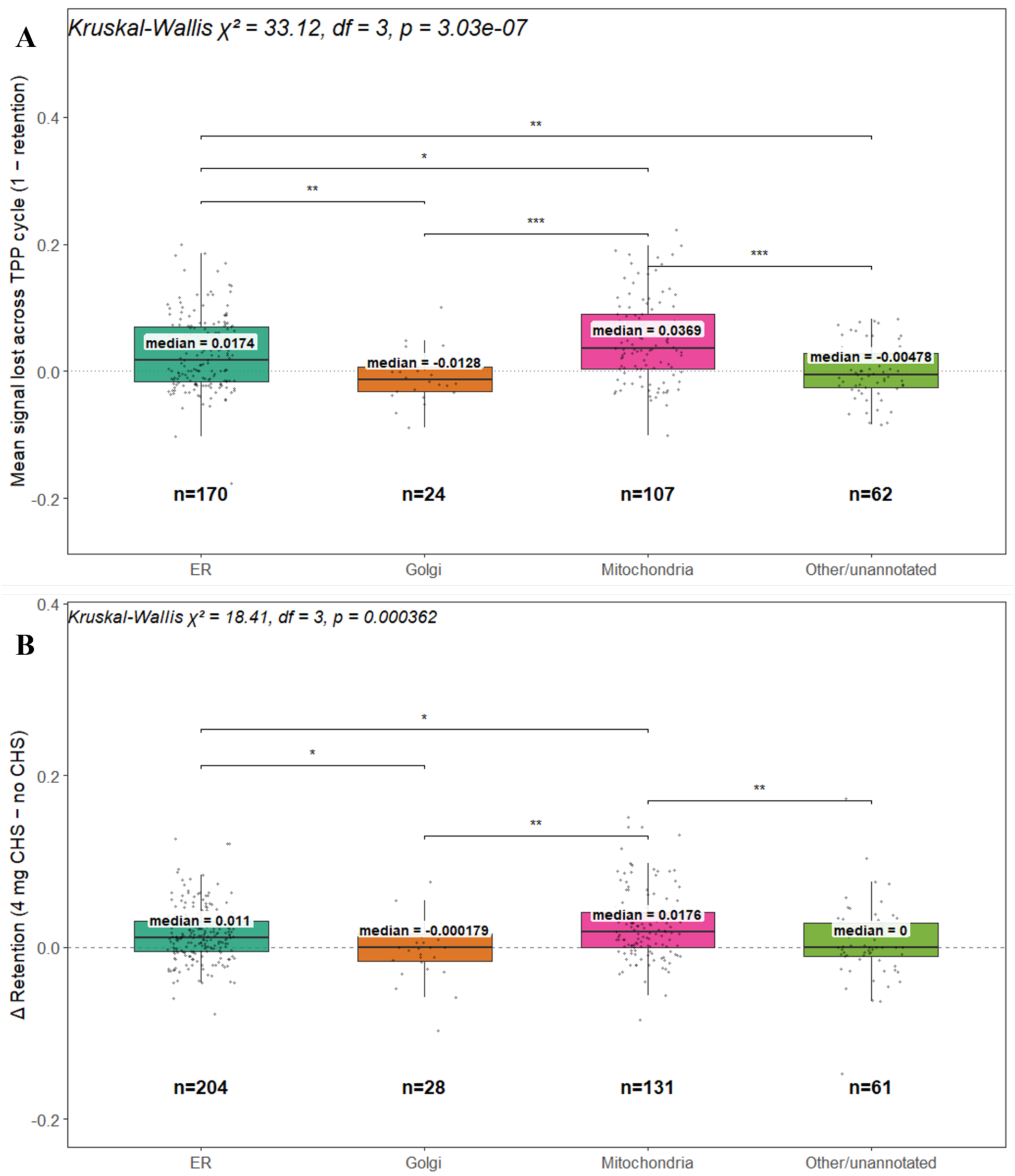
DDM solubilized oIMPs show distinct baseline thermal lability and differential CHS-induced stabilization across mitochondria, ER, Golgi, and other subcellular compartments. oIMP proteins detected in DDM were stratified by subcellular localization (GO cellular component) into ER, Golgi, Mitochondria, and Other/unannotated categories. **(A)** Baseline thermal lability, measured as mean signal lost across the TPP cycle (1 − retention) in the absence of CHS, differed significantly across organelle categories (Kruskal-Wallis χ² = 33.12, df = 3, p = 3.03×10⁻⁷). Mitochondrial proteins showed the greatest thermal lability (median = 0.0369), followed by ER (median = 0.0174), while Golgi (median = −0.0128) and Other/unannotated (median = −0.00478) proteins were comparatively thermally stable. Pairwise BH-adjusted Wilcoxon contrasts showed significant differences between Other/unannotated and Mitochondria (p = 1.25×10⁻⁵), Mitochondria and Golgi (p = 1.32×10⁻⁴), Golgi and ER (p = 3.03×10⁻³), Other/unannotated and ER (p = 3.03×10⁻³), and Mitochondria and ER (p = 0.0138); Other/unannotated and Golgi were not significantly different (p = 0.345). **(B)** CHS-induced retention shift (Δ retention = 4 mg CHS − no CHS) also differed significantly by organelle (Kruskal-Wallis χ² = 18.41, df = 3, p = 0.000362), with Mitochondria showing the largest median rescue (0.0176), followed by ER (0.011), while Golgi (−0.000179) and Other/unannotated (0) showed minimal shift. Pairwise BH-adjusted Wilcoxon contrasts showed significant differences between Mitochondria and Golgi (p = 0.00428), Other/unannotated and Mitochondria (p = 0.00501), Golgi and ER (p = 0.0342), and Mitochondria and ER (p = 0.0342); Other/unannotated and ER (p = 0.0619) and Other/unannotated and Golgi (p = 0.389) were not significant. Boxplots show median (labeled) and interquartile range; the dashed/dotted line marks zero; points represent individual proteins; n denotes protein number per category. Statistical analysis was performed in R using kruskal.test and pairwise.wilcox.test (Benjamini-Hochberg adjusted), with data processing in dplyr/tidyr and figures generated using ggplot2 and ggsignif.

**Supplemental Table 3.** Transmembrane helix count differs between pIMP and oIMP proteins in Peptidisc but not DDM. A Wilcoxon rank-sum test was used to compare transmembrane (TM) helix count between oIMP and pIMP proteins, independent of CHS-induced stabilization, to assess whether these localization classes differ structurally in TM topology. TM helix counts were derived from Phobius-predicted transmembrane topology annotations. In Peptidisc, pIMP proteins had significantly higher TM count than oIMP proteins (median = 2 vs. 1; mean = 4.36 vs. 2.52; W = 29934, p = 2.79×10⁻⁵, n = 553). In DDM, pIMP proteins showed a similar trend toward higher TM count (median = 1 vs. 1; mean = 3.80 vs. 2.81), but the difference did not reach statistical significance (W = 78841.5, p = 0.166, n = 840). All statistical analyses were performed in R using base stats functions. N refers to the number of oIMP and pIMP proteins with an annotated TM segment count.

| Dataset | Analysis | Test | Statistic | P-value | N |
| --- | --- | --- | --- | --- | --- |
| Peptidisc | TM count, pIMP vs. oIMP | Wilcoxon rank-sum | W=29934 | 2.79e-05 | 553 |
| DDM | TM count, pIMP vs. oIMP | Wilcoxon rank-sum | W=78841.5 | 0.166 | 840 |

**Supplemental Figure 6.**
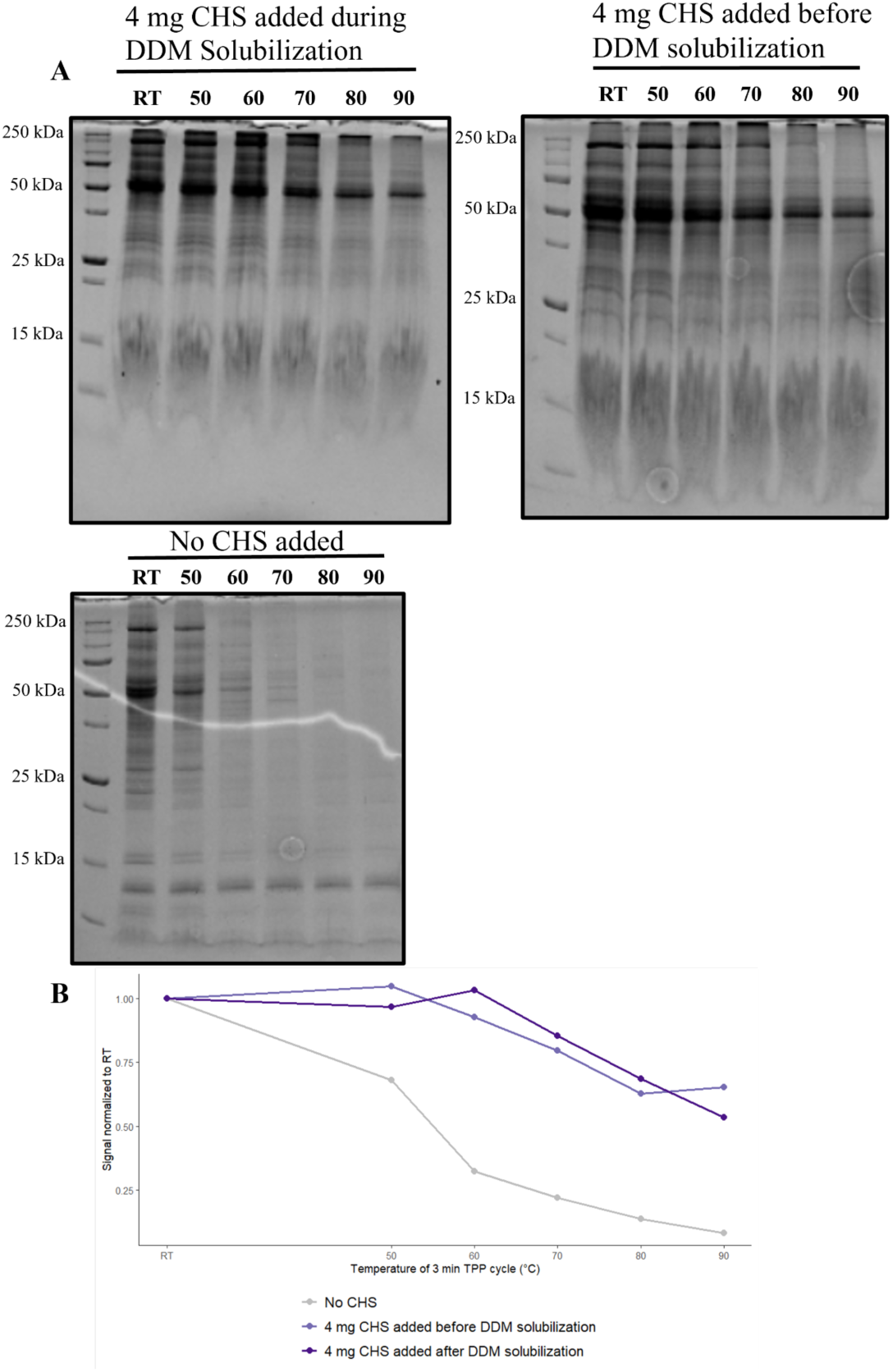
Effect of CHS Supplementation Before or During DDM Solubilization on Membrane Proteome Thermal Profiles. The mouse liver membrane proteome was solubilized in DDM under three conditions: no CHS added, 4 mg CHS added during DDM solubilization, or 4 mg CHS added before DDM solubilization. All preparations were subjected to TPP by 3-minute heat treatment at the indicated temperatures (RT–90°C), followed by ultracentrifugation to pellet aggregated proteins. **(A)** SDS-PAGE of the soluble supernatant fractions across the TPP temperature gradient for each condition. Molecular weight markers are indicated on the left (kDa). **(B)** Densitometric quantification of whole-lane signal from the SDS-PAGE gels shown in (A), normalized to the room temperature (RT) lane value for each condition. Signal intensities at 23°C were used as the RT reference, consistent with the x-axis labeling in the code. Densitometry was performed using ImageJ, and retention curves were plotted using ggplot2.

**Supplemental Figure 7.**
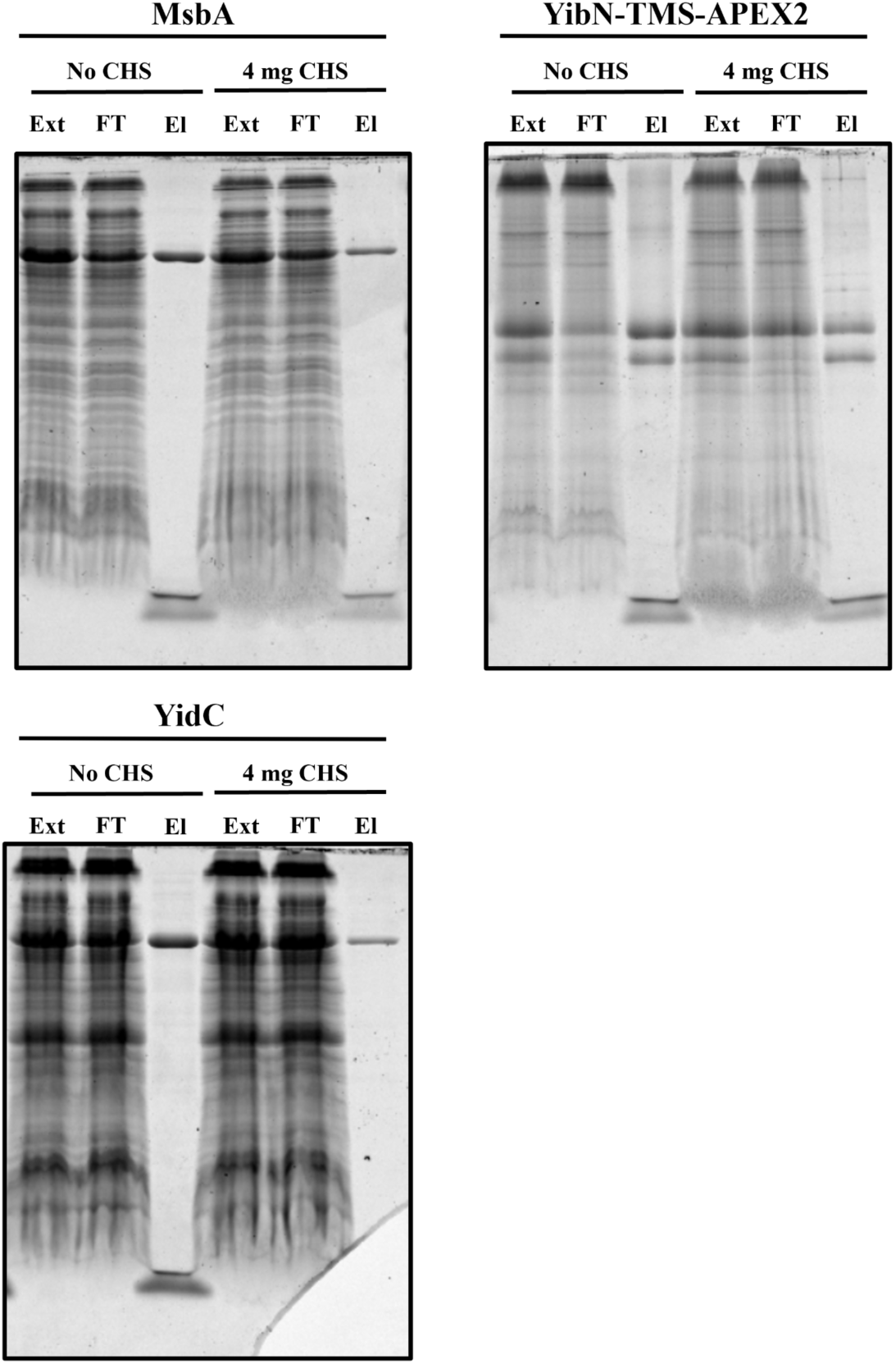
CHS supplementation during solubilization reduces elution yield of MsbA, YibN-TMS-APEX2, and YidC following Peptidisc reconstitution. MsbA, YibN-TMS-APEX2, and YidC, each bearing an N-terminal 6×His tag, were recombinantly expressed in *E. coli* and membrane fractions were solubilized in DDM with or without 4 mg CHS. Following solubilization, proteins were reconstituted into Peptidisc and purified by Ni-NTA affinity chromatography. Equal volumes of the cell extract (Ext), flow-through (FT), and elution (El) fractions were resolved by SDS-PAGE and stained with Coomassie Blue. Molecular weight markers are indicated in kDa.

**Supplemental Figure 8.**
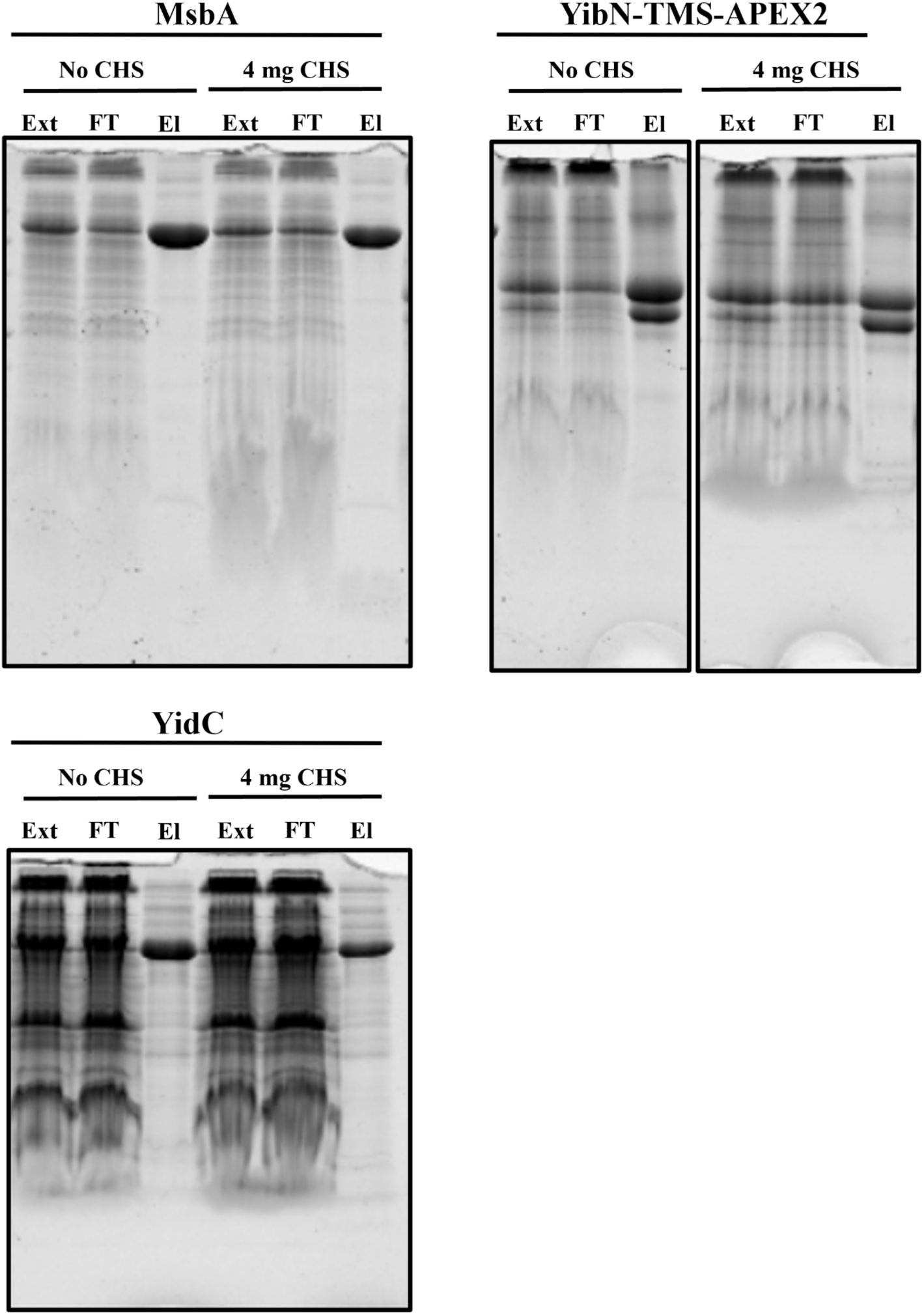
DDM MsbA, YibN-TMS-APEX2, and YidC are purified to comparable yield and purity with or without CHS supplementation during solubilization. MsbA, YibN-TMS-APEX2, and YidC, each bearing an N-terminal 6×His tag, were recombinantly expressed in *E. coli* and membrane fractions were solubilized in DDM with or without 4 mg CHS. Proteins were purified by Ni-NTA affinity chromatography, and equal volumes of the cell extract (Ext), flow-through (FT), and elution (El) fractions were resolved by SDS-PAGE and stained with Coomassie Blue. Molecular weight markers are indicated in kDa.

**Supplemental Figure 9.**
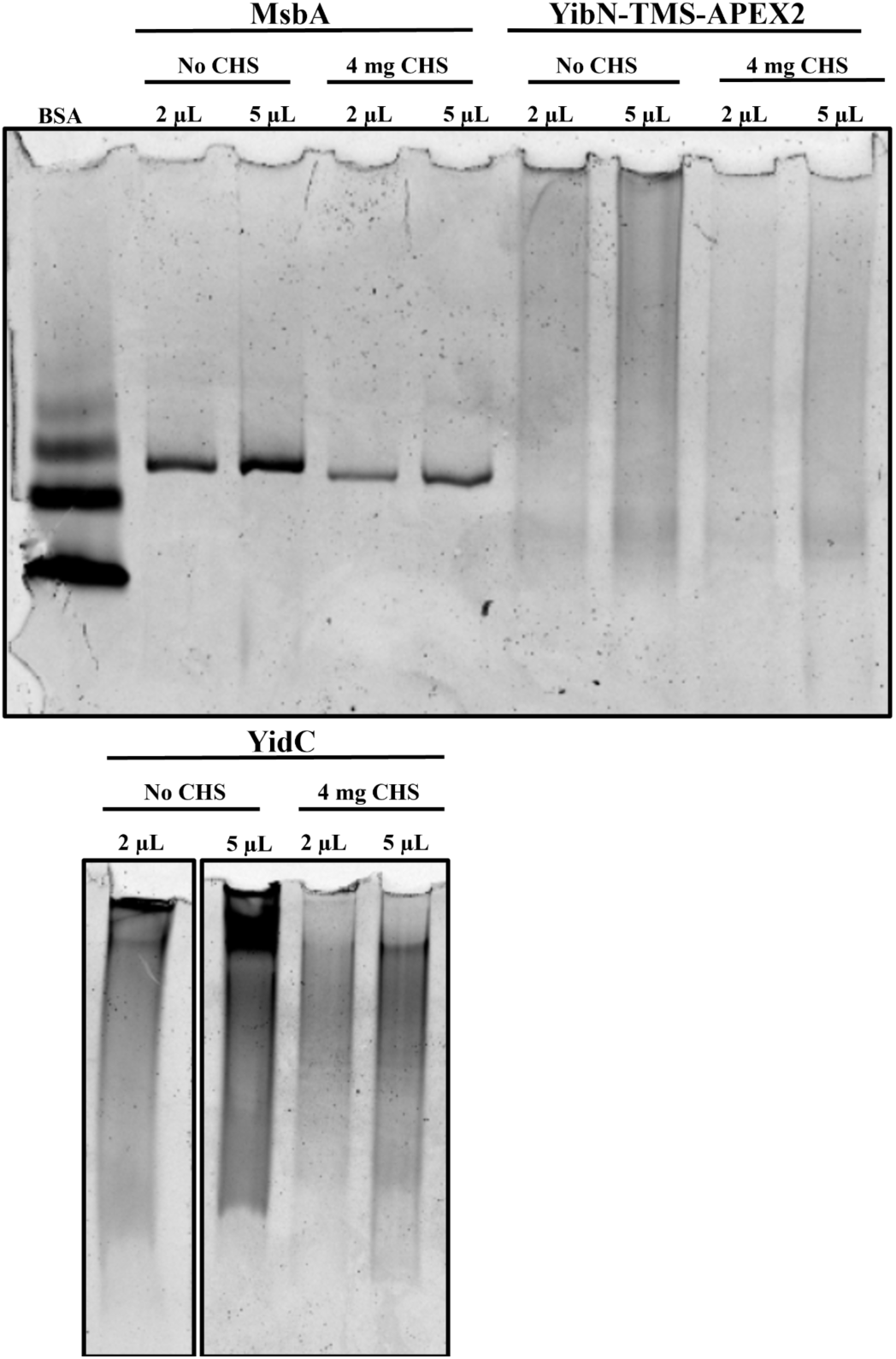
Clear Native PAGE shows CHS does not alter the oligomeric state of purified Peptidisc MsbA, YibN-TMS-APEX2, or YidC. Peptidisc reconstituted and purified MsbA, YibN-TMS-APEX2, and YidC were analyzed by Clear Native PAGE to assess native molecular weight and oligomeric state in a detergent-free membrane mimetic environment. Two sample volumes (2 µL and 5 µL) were loaded per condition to facilitate band detection across a range of protein concentrations. BSA was used as a molecular weight reference standard.

**Supplemental Figure 10.**
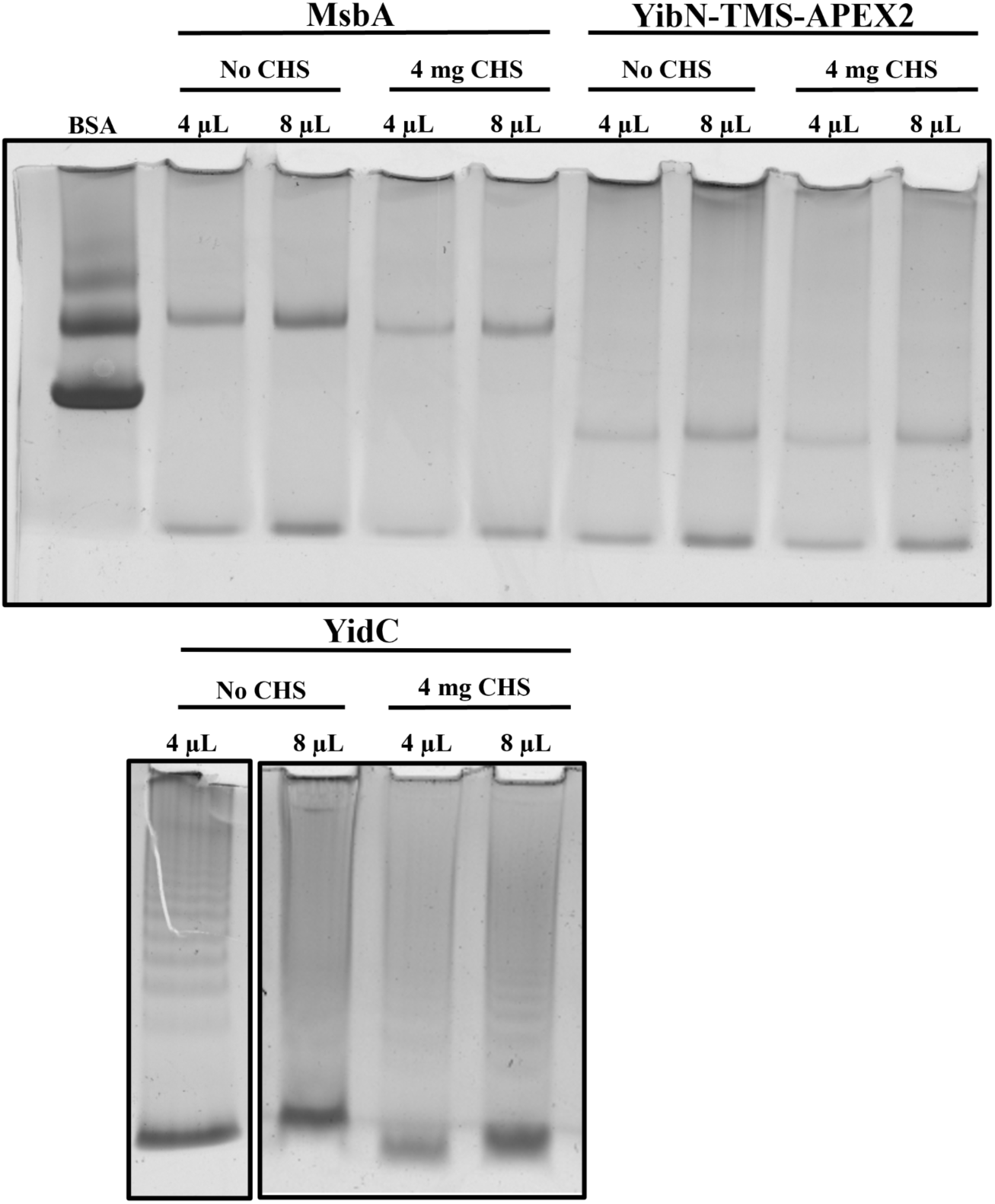
Blue Native PAGE shows CHS does not alter the oligomeric state of purified Peptidisc MsbA, YibN-TMS-APEX2, or YidC. Peptidisc reconstituted and purified MsbA, YibN-TMS-APEX2, and YidC were analyzed by Blue Native PAGE to assess native molecular weight and oligomeric state. Two sample volumes (4 µL and 8 µL) were loaded per condition. BSA was used as a molecular weight reference standard.

**Supplemental Figure 11.**
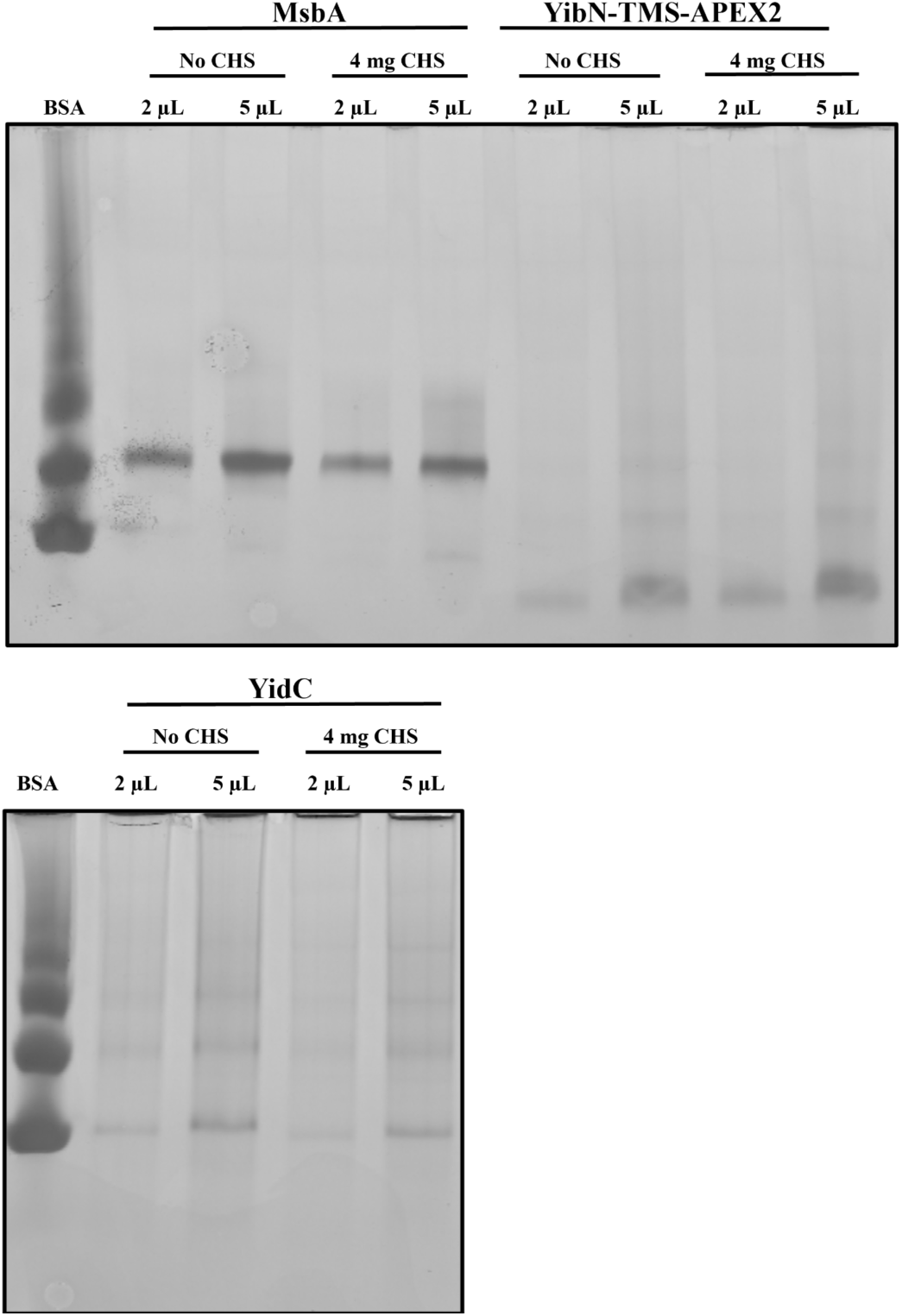
Blue Native PAGE shows CHS does not alter the oligomeric state of purified DDM MsbA, YibN-TMS-APEX2, or YidC. DDM solubilized and purified MsbA, YibN-TMS-APEX2, and YidC were analyzed by Blue Native PAGE to assess native molecular weight and oligomeric state. Two sample volumes (2 µL and 5 µL) were loaded per condition to facilitate band detection across a range of protein concentrations. BSA was used as a molecular weight reference standard.

**Supplemental Figure 12.**
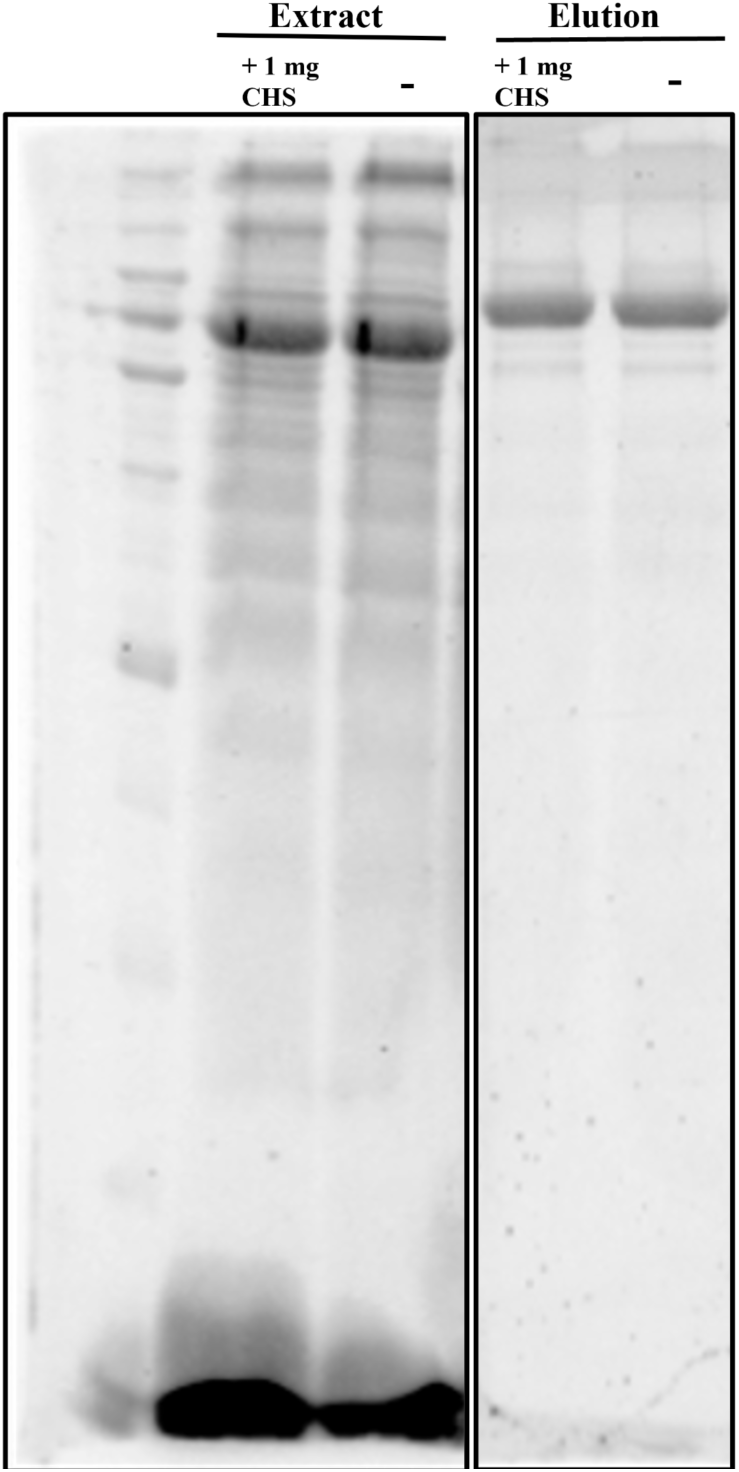
CHS does not alter the extraction or purification of MsbA in PDET-1. MsbA was solubilized using PDET-1 with or without 1 mg CHS. Crude membrane extract (Extract) and Ni-NTA imidazole elution (Ni-NTA Purified) fractions were resolved by SDS-PAGE and stained with Coomassie. Molecular weight markers are indicated on the left (kDa).

**Supplemental Figure 13.**
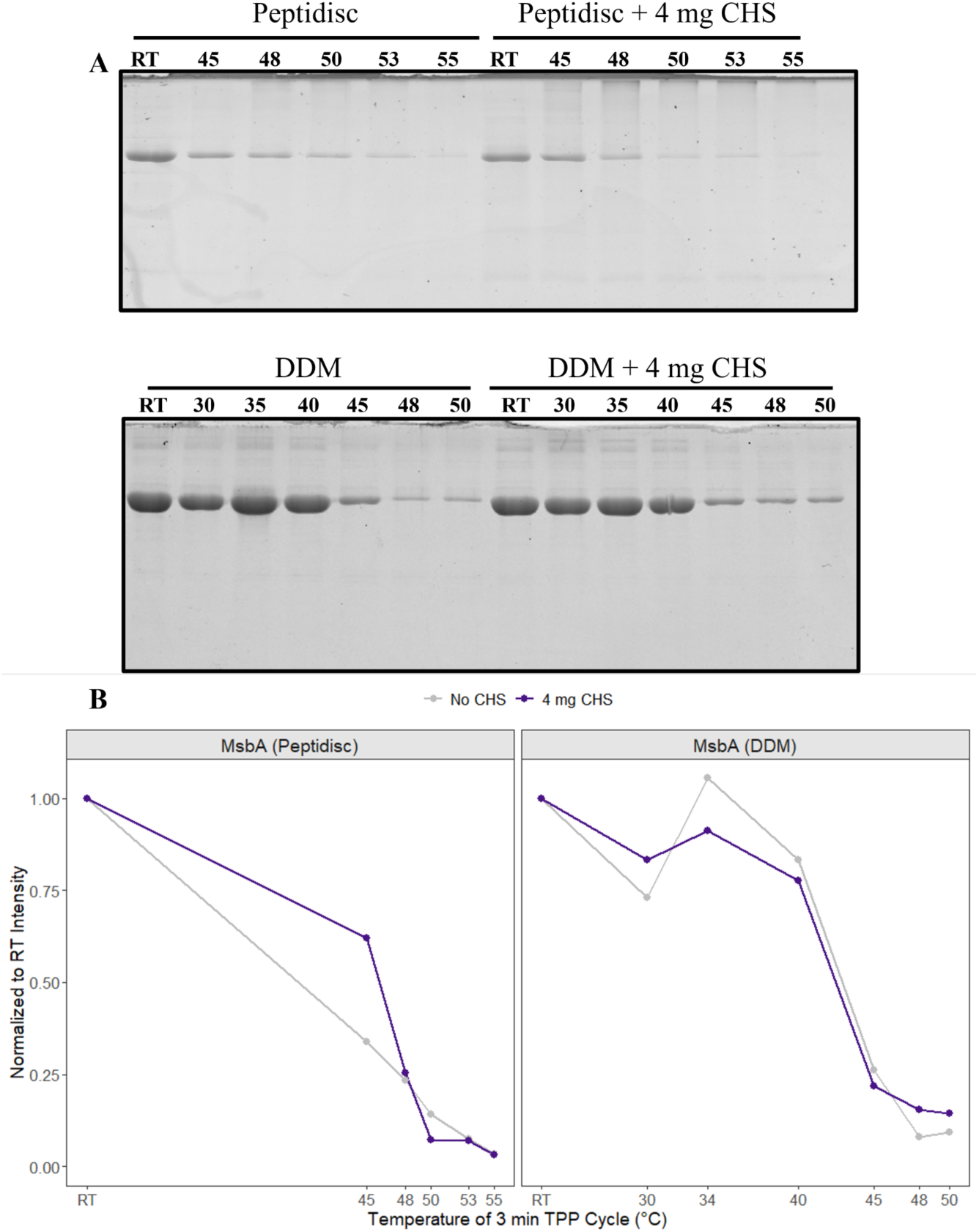
CHS-Associated Thermal Responses of Purified MsbA in DDM and Peptidisc. **(A)** Peptidisc reconstituted and DDM solubilized MsbA supplemented with and without CHS were subjected to 3-minute thermal treatment at the indicated temperatures, followed by ultracentrifugation to pellet aggregated protein, and the soluble supernatant was analyzed by SDS-PAGE. **(B)** Densitometric quantification of whole-lane signal from the SDS-PAGE gels shown in (A), normalized to the room temperature (RT) lane value for each condition performed with ImageJ. The two panels retain independent x-axes reflecting their different temperature ranges, rendered using facetted_pos_scales from ggh4x and plotted with ggplot2.

**Supplemental Figure 14.**
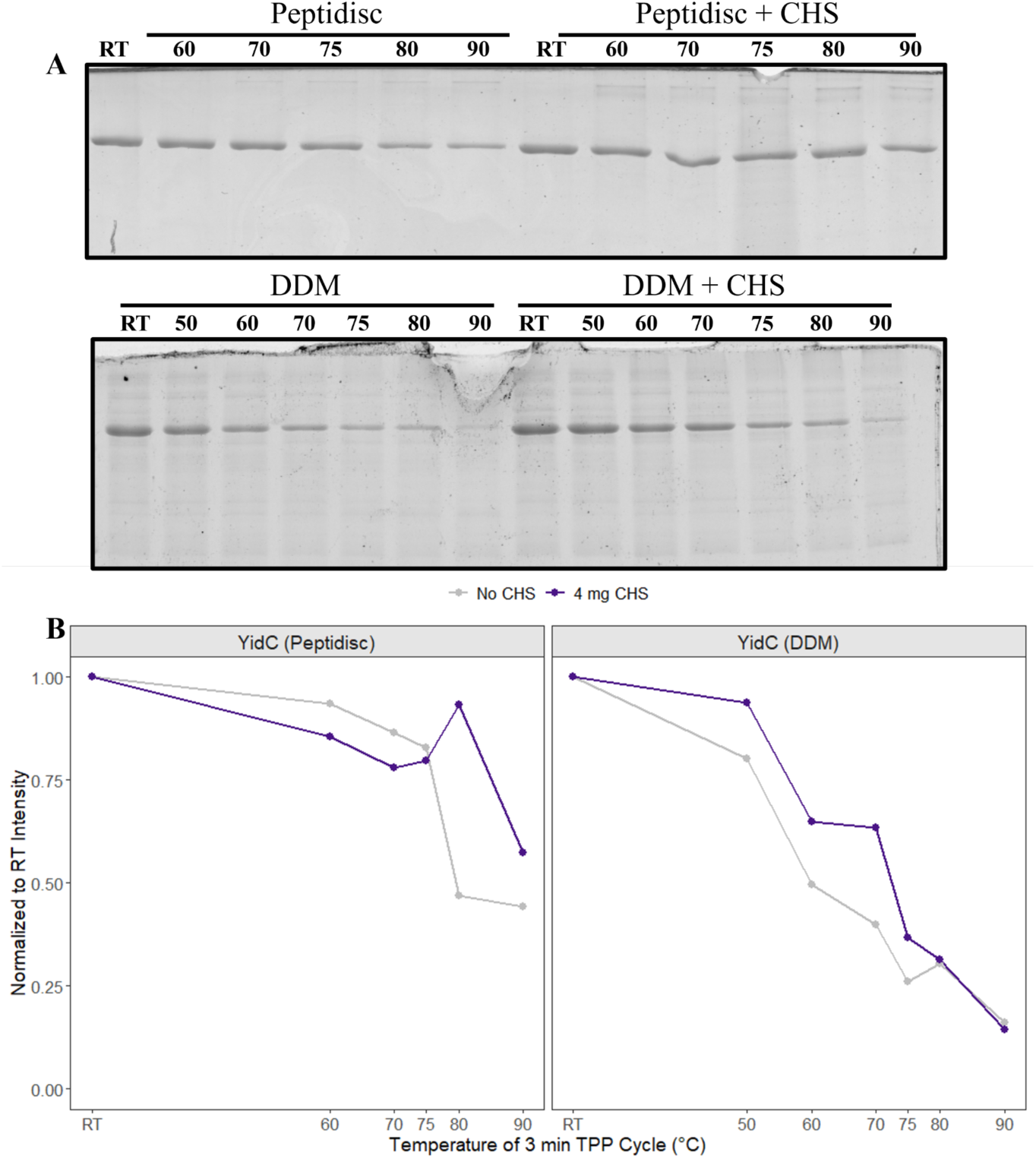
CHS-Associated Thermal Responses of Purified YidC in DDM and Peptidisc. **(A)** Peptidisc-reconstituted and DDM-solubilized YidC supplemented with or without 4 mg CHS were subjected to 3-minute thermal treatment at the indicated temperatures, followed by ultracentrifugation to pellet aggregated protein, and the soluble supernatant was analyzed by SDS-PAGE. Molecular weight markers are indicated in kDa. **(B)** Densitometric quantification of whole-lane signal from the SDS-PAGE gels shown in (A), normalized to the room temperature (RT) lane value for each condition performed with ImageJ. The two panels retain independent x-axes reflecting their different temperature ranges, rendered using facetted_pos_scales from ggh4x and plotted with ggplot2.

**Supplemental Table 4:**
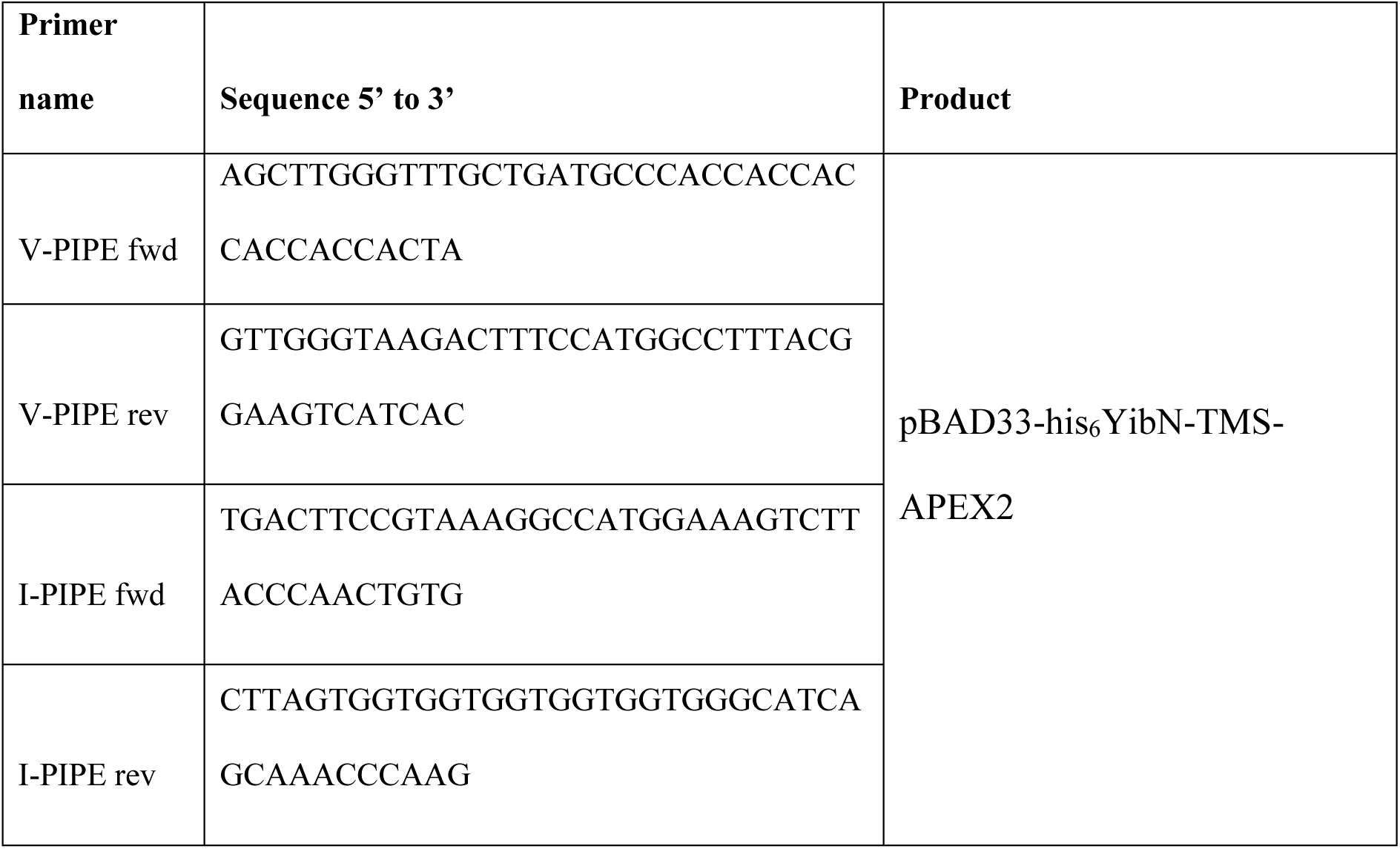
Primers used in the study.

## References

1. Das, A., Brown, M.S., Anderson, D.D., Goldstein, J.L., and Radhakrishnan, A. (2014). Three pools of plasma membrane cholesterol and their relation to cholesterol homeostasis. eLife 3, e02882. 10.7554/eLife.02882.

2. Ridsdale, A., Denis, M., Gougeon, P.-Y., Ngsee, J.K., Presley, J.F., and Zha, X. Cholesterol Is Required for Efficient Endoplasmic Reticulum-to-Golgi Transport of Secretory Membrane Proteins V□.

3. Shi, Q., Chen, J., Zou, X., and Tang, X. (2022). Intracellular Cholesterol Synthesis and Transport. Front. Cell Dev. Biol. 10, 819281. 10.3389/fcell.2022.819281.

4. Subczynski, W.K., Pasenkiewicz-Gierula, M., Widomska, J., Mainali, L., and Raguz, M. (2017). High Cholesterol/Low Cholesterol: Effects in Biological Membranes: A Review. Cell Biochem. Biophys. 75, 369–385. 10.1007/s12013-017-0792-7.

5. Zakany, F., Kovacs, T., Panyi, G., and Varga, Z. (2020). Direct and indirect cholesterol effects on membrane proteins with special focus on potassium channels. Biochim. Biophys. Acta BBA - Mol. Cell Biol. Lipids 1865, 158706. 10.1016/j.bbalip.2020.158706.

6. Bastiaanse, E. (1997). The effect of membrane cholesterol content on ion transport processes in plasma membranes. Cardiovasc. Res. 33, 272–283. 10.1016/S0008-6363(96)00193-9.

7. Scanlon, S.M., Williams, D.C., and Schloss, P. (2001). Membrane Cholesterol Modulates Serotonin Transporter Activity. Biochemistry 40, 10507–10513. 10.1021/bi010730z.

8. Brannigan G, Hénin J, Law R, Eckenhoff R, Klein ML. Embedded cholesterol in the nicotinic acetylcholine receptor. Proc Natl Acad Sci USA 2008;105(38):14418–14423. doi:10.1073/pnas.0803029105

9. Pöhnl, M., Trollmann, M.F.W., and Böckmann, R.A. (2023). Nonuniversal impact of cholesterol on membranes mobility, curvature sensing and elasticity. Nat. Commun. 14, 8038. 10.1038/s41467-023-43892-x.

10. Harayama, T., and Riezman, H. (2018). Understanding the diversity of membrane lipid composition. Nat. Rev. Mol. Cell Biol. 19, 281–296. 10.1038/nrm.2017.138.

11. Bieberich, E. (2018). Sphingolipids and lipid rafts: Novel concepts and methods of analysis. Chem. Phys. Lipids 216, 114–131. 10.1016/j.chemphyslip.2018.08.003.

12. Davis, J.H., Clair, J.J., and Juhasz, J. (2009). Phase Equilibria in DOPC/DPPC-d62/Cholesterol Mixtures. Biophys. J. 96, 521–539. 10.1016/j.bpj.2008.09.042.

13. Grisshammer, R. (2009). Chapter 36 Purification of Recombinant G-Protein-Coupled Receptors. In Methods in Enzymology (Elsevier), pp. 631–645. 10.1016/S0076-6879(09)63036-6.

14. Naranjo, A.N., McNeely, P.M., Katsaras, J., and Robinson, A.S. (2016). Impact of purification conditions and history on A2A adenosine receptor activity: The role of CHAPS and lipids. Protein Expr. Purif. 124, 62–67. 10.1016/j.pep.2016.05.015.

15. Antony, F., Bhattacharya, A., Aoki, H., Jandu, R.S., Abdualkader, A.M., Al Batran, R., Babu, M., and Duong Van Hoa, F. (2026). Comparative Evaluation of Solid-phase and Membrane Mimetic Strategies in Membrane Proteome Coverage and Disease-State Analysis. Mol. Cell. Proteomics 25, 101496. 10.1016/j.mcpro.2025.101496.

16. Denisov, I.G., and Sligar, S.G. (2017). Nanodiscs in Membrane Biochemistry and Biophysics. Chem. Rev. 117, 4669–4713. 10.1021/acs.chemrev.6b00690.

17. Carlson, M.L., Young, J.W., Zhao, Z., Fabre, L., Jun, D., Li, J., Li, J., Dhupar, H.S., Wason, I., Mills, A.T., et al. (2018). The Peptidisc, a simple method for stabilizing membrane proteins in detergent-free solution. eLife 7, e34085. 10.7554/eLife.34085.

18. Zhao, Z., Khurana, A., Antony, F., Young, J.W., Hewton, K.G., Brough, Z., Zhong, T., Parker, S.J., and Duong Van Hoa, F. (2023). A Peptidisc-Based Survey of the Plasma Membrane Proteome of a Mammalian Cell. Mol. Cell. Proteomics 22, 100588. 10.1016/j.mcpro.2023.100588.

19. Brough, Z., Zhao, Z., and Duong Van Hoa, F. (2024). From bottom-up to cell surface proteomics: detergents or no detergents, that is the question. Biochem. Soc. Trans. 52, 1253–1263. 10.1042/BST20231020.

20. Antony, F., Brough, Z., Orangi, M., Al-Seragi, M., Aoki, H., Babu, M., and Duong Van Hoa, F. (2024). Sensitive Profiling of Mouse Liver Membrane Proteome Dysregulation Following a High-Fat and Alcohol Diet Treatment. PROTEOMICS 24, e202300599. 10.1002/pmic.202300599.

21. Bhattacharya, A., Antony, F., Aoki, H., Babu, M., Ferguson, S.S.G., Abd-Elrahman, K.S., and Duong Van Hoa, F. (2026). Membrane Proteome Remodeling in Female APP/PS1 Mice Following M1 Muscarinic Receptor Modulation Revealed by Peptidisc-Enabled DIA-MS. J. Proteome Res. 25, 3136– 3148. 10.1021/acs.jproteome.6c00140.

22. Jandu, R.S., Yu, H., Zhao, Z., Le, H.T., Kim, S., Huan, T., and Duong van Hoa, F. (2024). Capture of endogenous lipids in peptidiscs and effect on protein stability and activity. iScience 27, 109382. 10.1016/j.isci.2024.109382.

23. Jandu, R.S., Bhattacharya, A., Antony, F., Al-Seragi, M., Aoki, H., Babu, M., and Duong Van Hoa, F. (2025). Membrane-mimetic thermal proteome profiling (MM-TPP) toward mapping membrane protein–ligand dynamic interactions. eLife 14, RP104549. 10.7554/eLife.104549.

24. Savitski, M.M., Reinhard, F.B.M., Franken, H., Werner, T., Savitski, M.F., Eberhard, D., Molina, D.M., Jafari, R., Dovega, R.B., Klaeger, S., et al. (2014). Tracking cancer drugs in living cells by thermal profiling of the proteome. Science 346, 1255784. 10.1126/science.1255784.

25. Jaakola, V.-P., Griffith, M.T., Hanson, M.A., Cherezov, V., Chien, E.Y.T., Lane, J.R., IJzerman, A.P., and Stevens, R.C. (2008). The 2.6 Angstrom Crystal Structure of a Human A_2A_ Adenosine Receptor Bound to an Antagonist. Science 322, 1211–1217. 10.1126/science.1164772.

26. Liu, Y., Liu, A., Li, X., Liao, Q., Zhang, W., Zhu, L., and Ye, R.D. (2024). Cryo-EM structure of monomeric CXCL12-bound CXCR4 in the active state. Cell Rep. 43, 114578. 10.1016/j.celrep.2024.114578.

27. Baier, C.J., Fantini, J., and Barrantes, F.J. (2011). Disclosure of cholesterol recognition motifs in transmembrane domains of the human nicotinic acetylcholine receptor. Sci. Rep. 1, 69. 10.1038/srep00069.

28. Li, H., and Papadopoulos, V. Peripheral-Type Benzodiazepine Receptor Function in Cholesterol Transport. Identification of a Putative Cholesterol Recognition/Interaction Amino Acid Sequence and Consensus Pattern. 139.

29. Antony, F., Bhattacharya, A., and Duong Van Hoa, F. (2026). PEPTERGENT: A Peptide-Based Reagent for Detergent-Free Extraction of Membrane Proteins and Purification of Membrane Proteomes. BIO-Protoc. 16. 10.21769/BioProtoc.5700.

30. Cherezov, V., Rosenbaum, D.M., Hanson, M.A., Rasmussen, S.G.F., Thian, F.S., Kobilka, T.S., Choi, H.-J., Kuhn, P., Weis, W.I., Kobilka, B.K., et al. (2007). High-Resolution Crystal Structure of an Engineered Human β_2_ -Adrenergic G Protein–Coupled Receptor. Science 318, 1258–1265. 10.1126/science.1150577.

31. Zhou, Y., Cao, C., He, L., Wang, X., and Zhang, X.C. (2019). Crystal structure of dopamine receptor D4 bound to the subtype selective ligand, L745870. eLife 8, e48822. 10.7554/eLife.48822.

32. El Daibani, A., Paggi, J.M., Kim, K., Laloudakis, Y.D., Popov, P., Bernhard, S.M., Krumm, B.E., Olsen, R.H.J., Diberto, J., Carroll, F.I., et al. (2023). Molecular mechanism of biased signaling at the kappa opioid receptor. Nat. Commun. 14, 1338. 10.1038/s41467-023-37041-7.

33. Weierstall, U., James, D., Wang, C., White, T.A., Wang, D., Liu, W., Spence, J.C.H., Bruce Doak, R., Nelson, G., Fromme, P., et al. (2014). Lipidic cubic phase injector facilitates membrane protein serial femtosecond crystallography. Nat. Commun. 5, 3309. 10.1038/ncomms4309.

34. Kim, K., Che, T., Panova, O., DiBerto, J.F., Lyu, J., Krumm, B.E., Wacker, D., Robertson, M.J., Seven, A.B., Nichols, D.E., et al. (2020). Structure of a Hallucinogen-Activated Gq-Coupled 5-HT2A Serotonin Receptor. Cell 182, 1574–1588.e19. 10.1016/j.cell.2020.08.024.

35. McCorvy, J.D., Wacker, D., Wang, S., Agegnehu, B., Liu, J., Lansu, K., Tribo, A.R., Olsen, R.H.J., Che, T., Jin, J., et al. (2018). Structural determinants of 5-HT2B receptor activation and biased agonism. Nat. Struct. Mol. Biol. 25, 787–796. 10.1038/s41594-018-0116-7.

36. Warren, A.L., Zilberg, G., Abbassi, A., Abraham, A., Yang, S., and Wacker, D. (2025). Structural determinants of G protein subtype selectivity at the serotonin receptor 5-HT1A. Sci. Adv.

37. Cecchetti, C., Strauss, J., Stohrer, C., Naylor, C., Pryor, E., Hobbs, J., Tanley, S., Goldman, A., and Byrne, B. (2021). A novel high-throughput screen for identifying lipids that stabilise membrane proteins in detergent based solution. PLOS ONE 16, e0254118. 10.1371/journal.pone.0254118.

38. Thompson, A.A., Liu, J.J., Chun, E., Wacker, D., Wu, H., Cherezov, V., and Stevens, R.C. (2011). GPCR stabilization using the bicelle-like architecture of mixed sterol-detergent micelles. Methods 55, 310–317. 10.1016/j.ymeth.2011.10.011.

39. Gimpl, G., and Fahrenholz, F. (2001). The Oxytocin Receptor System: Structure, Function, and Regulation. Physiol. Rev. 81, 629–683. 10.1152/physrev.2001.81.2.629.

40. Rosenbaum, D.M., Cherezov, V., Hanson, M.A., Rasmussen, S.G.F., Thian, F.S., Kobilka, T.S., Choi, H.-J., Yao, X.-J., Weis, W.I., Stevens, R.C., et al. (2007). GPCR Engineering Yields High-Resolution Structural Insights into β_2_ -Adrenergic Receptor Function. Science 318, 1266–1273. 10.1126/science.1150609.

41. Ruprecht, J.J., Mielke, T., Vogel, R., Villa, C., and Schertler, G.F. (2004). Electron crystallography reveals the structure of metarhodopsin I. EMBO J. 23, 3609–3620. 10.1038/sj.emboj.7600374.

42. Huang, S.K., Almurad, O., Pejana, R.J., Morrison, Z.A., Pandey, A., Picard, L.-P., Nitz, M., Sljoka, A., and Prosser, R.S. (2022). Allosteric modulation of the adenosine A2A receptor by cholesterol. eLife 11, e73901. 10.7554/eLife.73901.

43. Van Aalst, E., and Wylie, B.J. (2021). Cholesterol Is a Dose-Dependent Positive Allosteric Modulator of CCR3 Ligand Affinity and G Protein Coupling. Front. Mol. Biosci. 8, 724603. 10.3389/fmolb.2021.724603.

44. Howell, S.C., Mittal, R., Huang, L., Travis, B., Breyer, R.M., and Sanders, C.R. (2010). CHOBIMALT: A Cholesterol-Based Detergent. Biochemistry 49, 9572–9583. 10.1021/bi101334j.

45. Nji, E., Chatzikyriakidou, Y., Landreh, M., and Drew, D. (2018). An engineered thermal-shift screen reveals specific lipid preferences of eukaryotic and prokaryotic membrane proteins. Nat. Commun. 9, 4253. 10.1038/s41467-018-06702-3.

46. Serdiuk, T., Manna, M., Zhang, C., Mari, S.A., Kulig, W., Pluhackova, K., Kobilka, B.K., Vattulainen, I., and Müller, D.J. (2022). A cholesterol analog stabilizes the human β_2_ -adrenergic receptor nonlinearly with temperature. Sci. Signal. 15, eabi7031. 10.1126/scisignal.abi7031.

47. Li, S., Luo, H., Lou, R., Tian, C., Miao, C., Xia, L., Pan, C., Duan, X., Dang, T., Li, H., et al. (2021). Multiregional profiling of the brain transmembrane proteome uncovers novel regulators of depression. Sci. Adv. 7, eabf0634. 10.1126/sciadv.abf0634.

48. Perez-Riverol, Y., Bandla, C., Kundu, D.J., Kamatchinathan, S., Bai, J., Hewapathirana, S., John, N.S., Prakash, A., Walzer, M., Wang, S., et al. (2025). The PRIDE database at 20 years: 2025 update. Nucleic Acids Res. 53, D543–D553. 10.1093/nar/gkae1011.

49. Zhao, Z., Yamamoto, N., Young, J.W., Solis, N., Fong, A., Al-Seragi, M., Kim, S., Aoki, H., Phanse, S., Le, H.-T., et al. (2025). YibN, a bona fide interactor of the bacterial YidC insertase with effects on membrane protein insertion and membrane lipid production. J. Biol. Chem. 301, 108395. 10.1016/j.jbc.2025.108395.

50. Doyle, S.A. ed. (2009). High Throughput Protein Expression and Purification: Methods and Protocols (Humana Press) 10.1007/978-1-59745-196-3.

51. Chen, Y., and Duong Van Hoa, F. (2025). Peptidisc-Assisted Hydrophobic Clustering Toward the Production of Multimeric and Multispecific Nanobody Proteins. Biochemistry 64, 655–665. 10.1021/acs.biochem.4c00793.

52. Yu, F., Deng, Y., and Nesvizhskii, A.I. (2024). MSFragger-DDA+ Enhances Peptide Identification Sensitivity with Full Isolation Window Search. Preprint at Bioinformatics, 10.1101/2024.10.12.618041 https://doi.org/10.1101/2024.10.12.618041.

53. Kong, A.T., Leprevost, F.V., Avtonomov, D.M., Mellacheruvu, D., and Nesvizhskii, A.I. (2017). MSFragger: ultrafast and comprehensive peptide identification in mass spectrometry–based proteomics. Nat. Methods 14, 513–520. 10.1038/nmeth.4256.

54. Demichev, V., Messner, C.B., Vernardis, S.I., Lilley, K.S., and Ralser, M. (2020). DIA-NN: neural networks and interference correction enable deep proteome coverage in high throughput. Nat. Methods 17, 41–44. 10.1038/s41592-019-0638-x.

55. The UniProt Consortium, Bateman, A., Martin, M.-J., Orchard, S., Magrane, M., Adesina, A., Ahmad, S., Bowler-Barnett, E.H., Bye-A-Jee, H., Carpentier, D., et al. (2025). UniProt: the Universal Protein Knowledgebase in 2025. Nucleic Acids Res. 53, D609–D617. 10.1093/nar/gkae1010.

56. Käll, L., Krogh, A., and Sonnhammer, E.L.L. (2004). A Combined Transmembrane Topology and Signal Peptide Prediction Method. J. Mol. Biol. 338, 1027–1036. 10.1016/j.jmb.2004.03.016.

